# The superficial tufted and mitral cell output neurons of the mouse olfactory bulb have dual roles in insulin sensing

**DOI:** 10.1101/2025.04.15.648999

**Authors:** Louis John Kolling, Saptarsi Mitra, Catherine Anne Marcinkiewcz, Debra Ann Fadool

## Abstract

The olfactory bulb (OB) contains multiple, parallel projection neurons to relay the nature of a stimulus. In a mouse *ex vivo* slice preparation, we used patch-clamp electrophysiology to measure intrinsic properties, excitability, action potential (AP) shape, voltage-activated conductances, and neuromodulation in the newly-categorized superficial tufted cells (sTCs) compared with those of mitral cells (MCs). We propose that a marked difference in voltage-dependent current represents distinct ion channel populations that affect the kinetics of action potentials, and evokes an increase in sTC firing frequency, albeit both types of projection neurons having similar AP spiking activity. Triple-colored immunofluorescence and RNA scope were used to detect co-localization of the Kv1.3 ion channel and the insulin receptor in sTCs, with ∼73% of sTCs expressing both. The sTCs were modulated by bath application of insulin – increasing AP firing frequency by 97%, attributable to an 8% decrease in the intraburst interval, and a reduction of the latency to first spike by 37%. We conclude that there may be a range of neuromodulators of sTCs that may alter excitability and fine-tune olfactory information processing or metabolic balance.

**SUMMARY STATEMENT:** Superficial tufted cells, as output neurons of the olfactory bulb, were electrophysiologically studied to be insulin sensitive. Brain insulin signaling represents a manner in which olfactory and metabolic circuitry are intertwined.

## INTRODUCTION

Sensory systems typically carry multiple types of parallel output, or projection neurons, to relay differing specificity with regards to the nature of the stimulus. The olfactory bulb (OB) processes electrically-encoded odor information, and relays it to overlapping, but not identical, cortical regions via two types of primary projection neurons – namely mitral (MCs) and tufted cells (TCs) (Price and Powell, 1970; Vaaga and Westbrook, 2016). Albeit both receiving synaptic connections from the olfactory sensory neurons (OSNs) at defined and evolutionarily-conserved integration units called glomeruli, less has been studied concerning TCs in terms of biophysical properties and sources of modulation (Liu and Shipley, 2008; Burton and Urban, 2014; Jones et al., 2020). We and other investigators typically recorded from both populations of neurons, referring to them collectively as M/TCs (Fadool and Levitan, 1998; Tucker and Fadool, 2002; Youngstrom and Strowbridge, 2015; Liu et al., 2016). In model organisms, such as drosophila and zebrafish, MCs and TCs are not readily distinguishable from one another (Nagayama et al., 2014). The neurolaminae of mice are significantly more distinct, allowing the complexity of the olfactory circuity to be more easily assessed.

Like many budding areas of research, the standardization of categorical terminology is not considered until there is enough information to classify. Jones et al. 2020 aptly stated that the *“nomenclature around TC subtypes has been somewhat confusing in the field* (Jones et al., 2020).” Both early works and more recent reviews describe 4 distinct classes of projection neurons – external TC, internal TC, middle TC, and MC (Pinching and Powell, 1971a,b; Mori, 1987; Shepherd, 2004; Hayar et al., 2004; Mori and Sakano, 2022). A more recent work proposed to alternatively reclassify output neurons based upon size and location into three types: putative MCs (pMC; >20 um diameter and located in the MCL), EPL located TC (lower EPL TC; <20 um diameter), and putative TC (pTC; <20 um diameter located in the MCL) (Gadiwalla et al., 2025). Herein, our study uses the naming convention originating from Cajal and then reestablished by Jones et al. (2020), based upon the position of the TC soma within its respective neurolamina – namely, external tufted cells (eTCs), middle tufted cells (mTCs), and superficial tufted cells (sTCs) (**Fig. 1**). Of the three subclasses of TCs, the eTC are the best studied. The eTCs are a select population of juxtaglomerular cells that demonstrate rhythmic and spontaneous action potential bursting (Hayar et al., 2004a,b, 2005; Antal et al., 2006; Zhou et al., 2006; Ma and Lowe, 2007; Liu and Shipley, 2008). The eTC are known to receive monosynaptic input from olfactory sensory neurons (OSNs) and establish glutaminergic synaptic connections to short-axon cells and periglomerular cells within the bulb (Hayar et al., 2004a,b, 2005), and their activation can provide feedforward excitation of MC (De Saint Jan et al. 2009). eTCs actually process sensory input before it reaches the sTCs and MCs (De Saint Jan et al., 2009; Najac et al., 2011; Gire et al., 2012; Igarashi et al., 2012; Vaaga and Westbrook, 2016). The eTCs receive the greatest signal from activated OSNs, and stimulate neighboring sTCs, mTCs, and MCs via glutamatergic dendro-dendritic interactions (De Saint Jan *et al*., 2009; Najac *et al*., 2011; Jones *et al*., 2020). This feed-forward excitation from eTCs provides “biphasic input” to other M/TCs that may also receive direct monosynaptic input from a specific OSN family member (Najac *et al*., 2011; Vaaga and Westbrook, 2016; Jones *et al*., 2020).

**Fig. 1.**
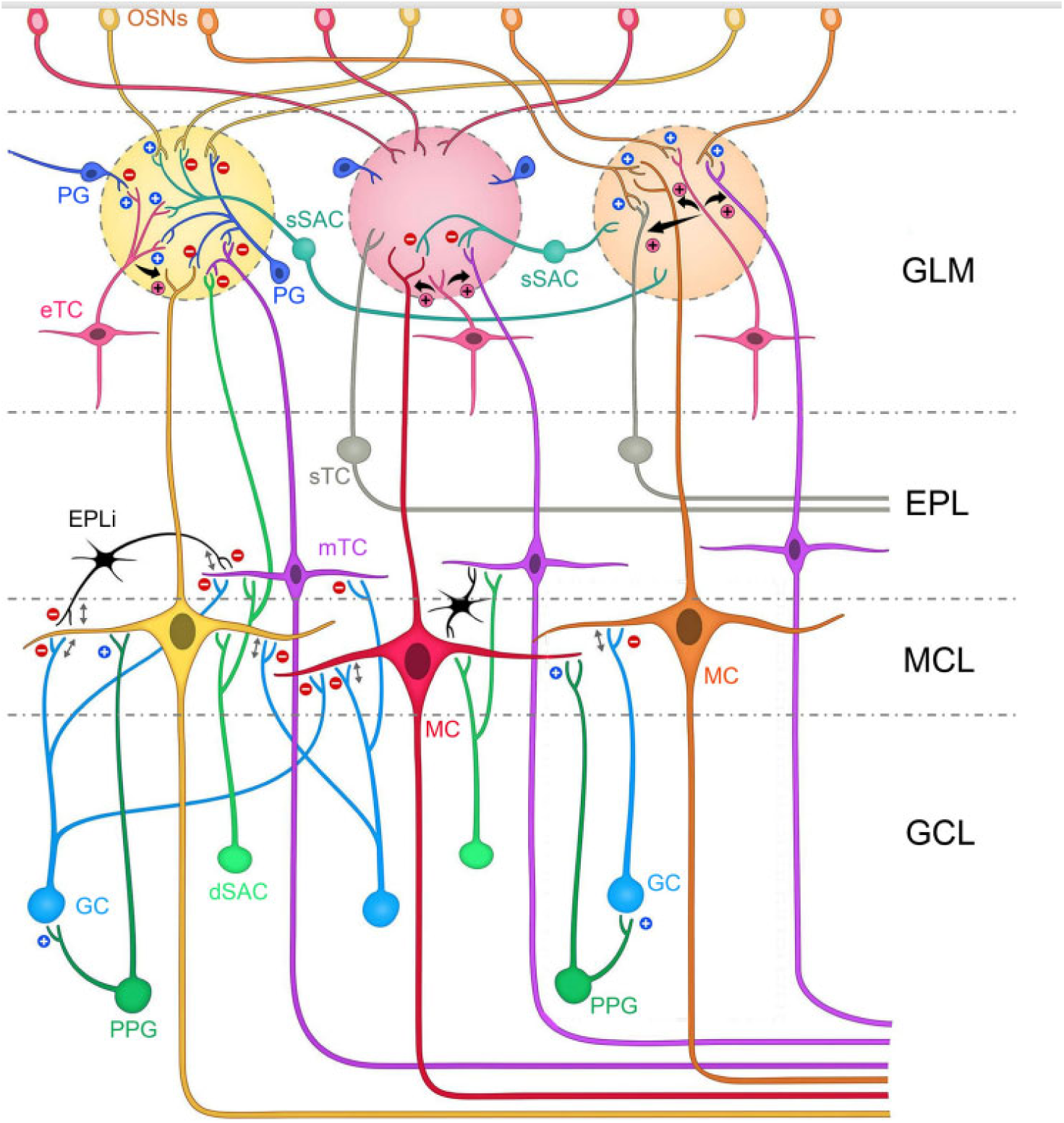
Categorization and interactions of the principal projection neurons of the olfactory bulb. The olfactory bulb neurolamina is divided across the glomerular layer (GLM), external plexiform layer (EPL), the mitral cell layer (MCL), and the innermost granule cell layer (GCL). Also shown is the olfactory mucosa containing the olfactory sensory neurons (OSNs). This schematic highlights the four classes of output or projection neurons, namely the external tufted cells (eTCs, pink), the superficial tufted cells (sTCs, gray), the middle tufted cells (mTCs, purple), and the mitral cells (MCs, red). The soma of the eTCs are found within the GLM. The sTC soma reside in the superficial one-third of the EPL. The deep two-thirds of the EPL contain both the mTCs and misplaced MCs. The MCL houses the eponymous MCs. Blue plus signs (+) represent excitatory synapses, red minus signs (-) represent inhibitory synapses, and pink plus signs (+) represent excitatory dendro-dendritic interactions originating from external tufted cells (eTCs). Reciprocal synapses are denoted by a double arrow. PG = periglomerular cell, sSAC = superficial short axon cell, EPLi = interneuron of the external plexiform layer, GCL = granule cell layer, GC = granule cell, PPG = preproglucagon neuron, dSAC = deep short axon cell. *Modified from Huang* et al. *2021*.

Because the eTC are well characterized, and the mTCs are less accessible for patch-clamp recordings, we decided to focus our study on a lesser known type of TC, the sTC, which are expressed superficially in the external plexiform layer (EPL)(Jones et al., 2020; Chen et al., 2024). mTC soma are found between the MCL and the inner (deep) two-thirds of the external plexiform layer (EPL), whereas the soma of superficial tufted cells (sTCs) are located in the outer (superficial) one-third of the EPL. Like MCs, sTCs and mTCs receive input from olfactory sensory neurons (OSNs) at defined glomeruli, and pass sensory information to various subcortical areas. The sTCs and mTCs, however, project, to distinctly different brain regions than MCs; even if a M/TC pair projects to the same subcortical region, it will project to different parts of it (Nagayama et al., 2010, 2014; Igarashi et al., 2012). Excellent foundational work has been done to characterize the innate biophysical differences between TCs and MCs (Burton and Urban, 2014), but the categorization of the assessed cells preceded the boundaries set by the current naming methods (Jones et al., 2020). As such, no previous report has characterized the innate biophysical differences between MCs and sTCs, nor investigated the function of sTCs to sense metabolically-relevant molecules or neuromodulators.

In this study, we compare the biophysical properties of MCs and sTCs in *ex vivo* brain slices from male and female mice. Using current- and voltage-clamp recordings, we found that action potential (AP) properties and voltage-activated currents differed between sTCs and MCs. To better understand the complementary function of sTCs to MCs, we examined the response of sTCs to a known modulator and a vestibule blocker of Kv1.3 – a predominant ion channel carrying 60-80% of the outward current in MCs (Fadool and Levitan, 1998; Colley et al., 2004). Our results directly show that sTCs are more excitable than MCs. This is correlated with differing voltage-activated currents, and a differing AP shape. We further find that sTCs are modulated by the presence of insulin, suggesting a role for sTCs in metabolic circuitry (Fadool et al., 2004; Palouzier-Paulignan et al., 2012; Aimé et al., 2014). While both MC and sTCs carry the role of propagating odor-coding information to higher cortical regions, they have another role, hence a dual role, to encode metabolic state when insulin is present, either locally or as modulated following food ingestion. The major olfactory output neurons hence have a dual role – one for odor encoding, and a second for changed activity with metabolism. Our study demonstrates that one downstream target, the Kv1.3 ion channel, is necessary for the insulin-sensing property of sTCs, as it is in MCs (Fadool et al., 2004, 2011; Tucker et al., 2013).

## RESULTS

### Superficial tufted cells have greater input resistance and slightly lower voltage-activated current at hyperpolarizing potentials than that of mitral cells

Superficial tufted cells (sTCs) have greater EPSC amplitude when stimulated by direct olfactory sensory neuron activation over that of mitral cells (MCs) (Jones et al., 2020), albeit both being the predominant output neurons of the OB. Less has been studied to reveal the nature of intrinsic properties and voltage-activated conductances in sTCs. We used ex vivo slice electrophysiology in the voltage-clamp configuration, therefore, to study intrinsic properties as well as outward currents, which are largely driven by the Kv1.3 channel in MCs (Fadool *et al*., 1997; Fadool and Levitan, 1998). The resting membrane potential (RMP) was not significantly different between MCs and sTCs (MCs = -55.8 ± 6.3 mV (20) vs. sTCs = - 58.0 ± 6.5 mV (22); Student’s *t*-test, *p* = 0.2906).

Macroscopic currents were acquired in response to a series of 400 ms duration (P_d_) voltage steps that increased in increments of 20 mV from -100 mV to +40 mV. The resulting current-voltage (IV) relationship was plotted in **Fig. 2A** using the peak transient current value for each step normalized to the +40 mV step. It was noted that the amplitude of the outward current at hyperpolarizing potentials (a window between -60 to -20 mV) was significantly greater in MCs vs. that of sTCs (**Fig. 2B**). A two-way repeated measure, mixed factor analysis of variance (2-w RM mixed ANOVA) was applied using cell type and voltage as factors. There was a main effect of cell type (F (1, 22) = 9.430; *p* = **0.0056) as well as a voltage x cell type interaction (F (7, 154) = 5.542; *p* <0.0001). A Sidak’s post-hoc test showed that the evoked current was significantly greater for MCs at the -60 mV (*p* = 0.0202), -40 mV (*p* = 0.0049) and -20 mV (*p* = 0.0311) voltage steps. This difference only existed in this hyperpolarized window and was not observed once highly depolarized at the +40 mV step (**Fig. 2C**, MCs = 5180 ± 2540 pA (11) vs. sTCs = 3746 ± 1030 pA (13), Mann-Whitney U, *p* = 0.2066). We next measured the kinetics of inactivation and deactivation for the voltage-activated currents at a hyperpolarized (V_c_ = -40 mV) and depolarized (V_c_ = +40 mV) range, respectively. The kinetics of inactivation (τ_Inact_) did not differ between cell types (**Fig. 2D, Left**, MCs = 71.2 ± 19.0 ms (10) vs. sTCs = 57.0 ± 28.6 ms (13), Student’s *t*-test, *p* = 0.1894) at V_c_ = -40 mV, however the kinetics of deactivation (τ_Deact_) was significantly faster for sTCs at this command voltage (**Fig. 2D, Right**, MCs = 2.19 ± 0.67 ms (11) vs. sTCs = 1.47 ± 0.61 ms (13), Mann-Whitney U, \**p* = 0.0107). Neither time constant differed between cell types at a V_c_ = +40 mV step (**Fig. 2E**, τ_Inact_: MCs = 75.6 ± 30.9 ms (11) vs. sTCs = 158 ± 140 ms (13), Mann-Whitney U, *p* = 0.3031; τ_Deact_: MCs = 1.14 ± 0.41 ms (10) vs. sTCs = 1.12 ± 0.52 ms (13), Student’s *t*-test, *p* = 0.9043). Overall, although sTCs tend to have higher input resistance than MCs, the outward voltage-activated currents of sTCs are less responsive to changes in voltage.

**Fig. 2.**
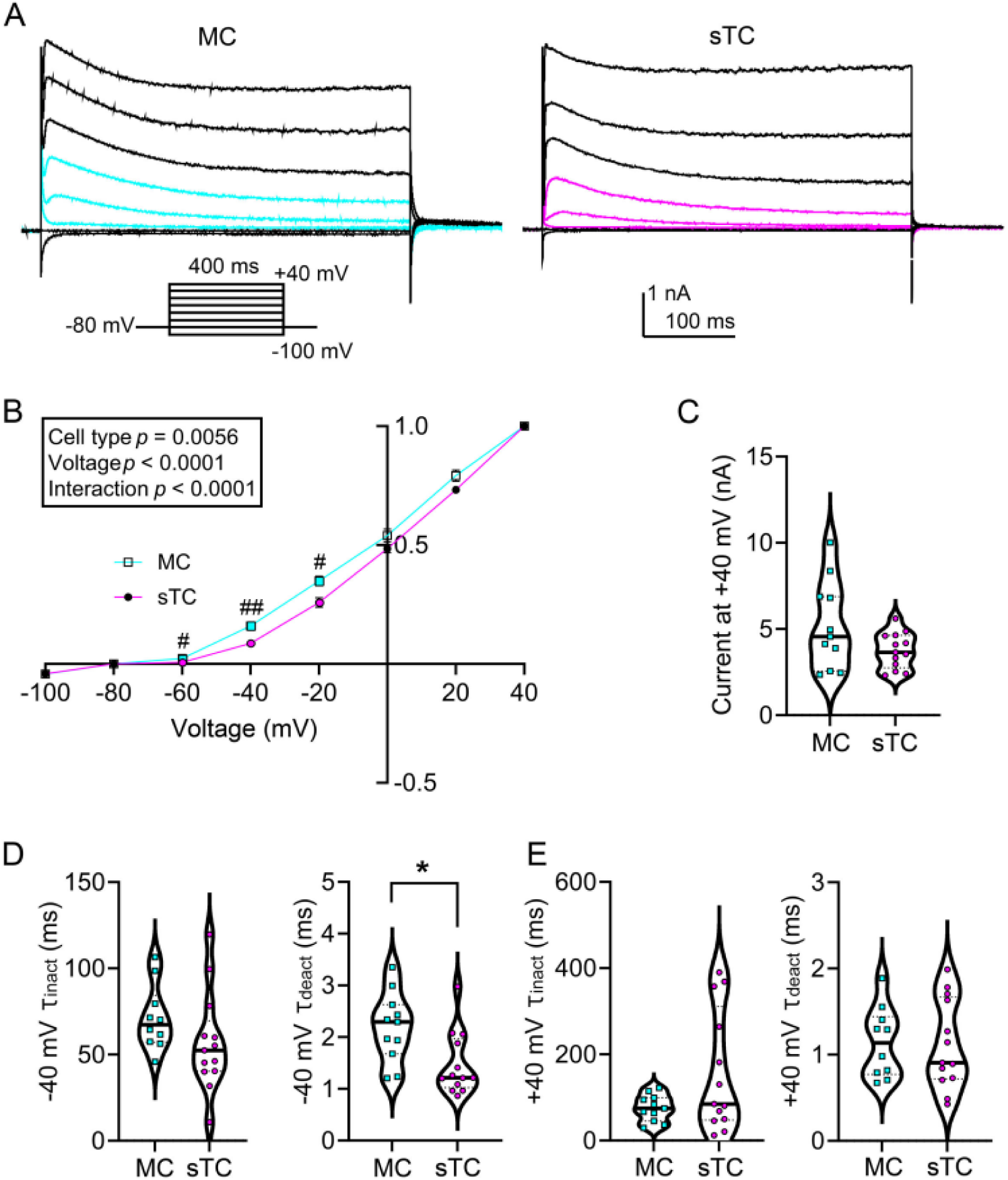
Outward voltage-activated current properties of mitral cells and superficial tufted cells. ***A***, Family of voltage-activated currents, V_h_ = -80 mV where V_c_ was stepped in 20 mV increments from - 100 mV to +40 mV using a P_d_ = 400 ms. Colored sweeps correspond to significantly different colored data points in (***B***). ***B***, Normalized IV plots (2-w mixed RM ANOVA, cell type \*\*\**p* = 0.0056, Sidak’s post-hoc test, ^#^*p* < 0.05, ^##^*p* < 0.01; voltage x cell type interaction \*\*\*\**p* < 0.0001). ***C***, Violin plot of non-normalized peak transient voltage-activated current at +40 mV, Mann-Whitney U. Violin plot of the *Tau* of inactivation (*left*) and deactivation (*right*) at **(D)** Vc = -40 mV and **(E)** +40 mV, respectively, Mann-Whitney U (\**p* = 0.0107). MC = mitral cell population sampled, sTC = superficial tufted cell population sampled.

### Kv1.2 and Kv1.3 expression is similar between superficial tufted and mitral cells

The drug MgTx is a selective blocker of Kv1.2 and Kv1.3 ion channels when applied to the extracellular membrane at 1 nM concentration (Garcia-Calvo et al., 1993; Garcia et al., 1997; Bartok et al., 2014). It has a slow *k*_on_ of 10 min and an even slower *k*_off_ of 2 h (Garcia-Calvo et al., 1993; Nikouee et al., 2012; Chen and Chung, 2014), therefore, we did not attempt a washout of the vestibule blocker during our patch recording. After collecting baseline voltage-clamp recordings on MCs for 15 min, 1 nM MgTx was added to the bath to wash over the slice for 15 min. Another set of voltage-clamp recordings was then collected (**Fig. 3A**). The same repeated-measure paradigm was then applied to a separate group of sTCs (**Fig. 3B**). MgTx reduced voltage-activated currents in both MCs and sTCs in a voltage-dependent manner (**Fig. 3C).** Here, a 2-w mixed RM ANOVA was applied using drug and voltage as factors. For the MC recordings, there was a main effect of drug (F (1, 9) = 43.62, \*\*\*\**p* < 0.0001) as well as a voltage x drug interaction (F (7, 63) = 43.86, \*\*\*\**p* < 0.0001). A Sidak’s post-hoc test demonstrated significant differences at the -20 mv (^###^*p* = 0.0008) voltage step and all further depolarizing steps (^####^*p* < 0.0001). For the sTC recordings, there was a main effect of drug (F (1, 12) = 25.51, \*\*\**p* = 0.0003) as well as a voltage x drug interaction (F (7, 60) = 19.87, \*\*\*\**p* < 0.0001). A Sidak’s post-hoc test demonstrated significant differences at -30 mv and above (^####^*p* < 0.0001). Using a 3-way mixed-effects analysis (3-w mixed RM ANOVA), we found no difference in MgTx-mediated effects on voltage dependence between MCs and sTCs (**Fig. 3C, Bottom**, voltage x cell type x drug interaction F (7, 123) = 0.7633, *p* = 0.6190).

**Fig. 3.**
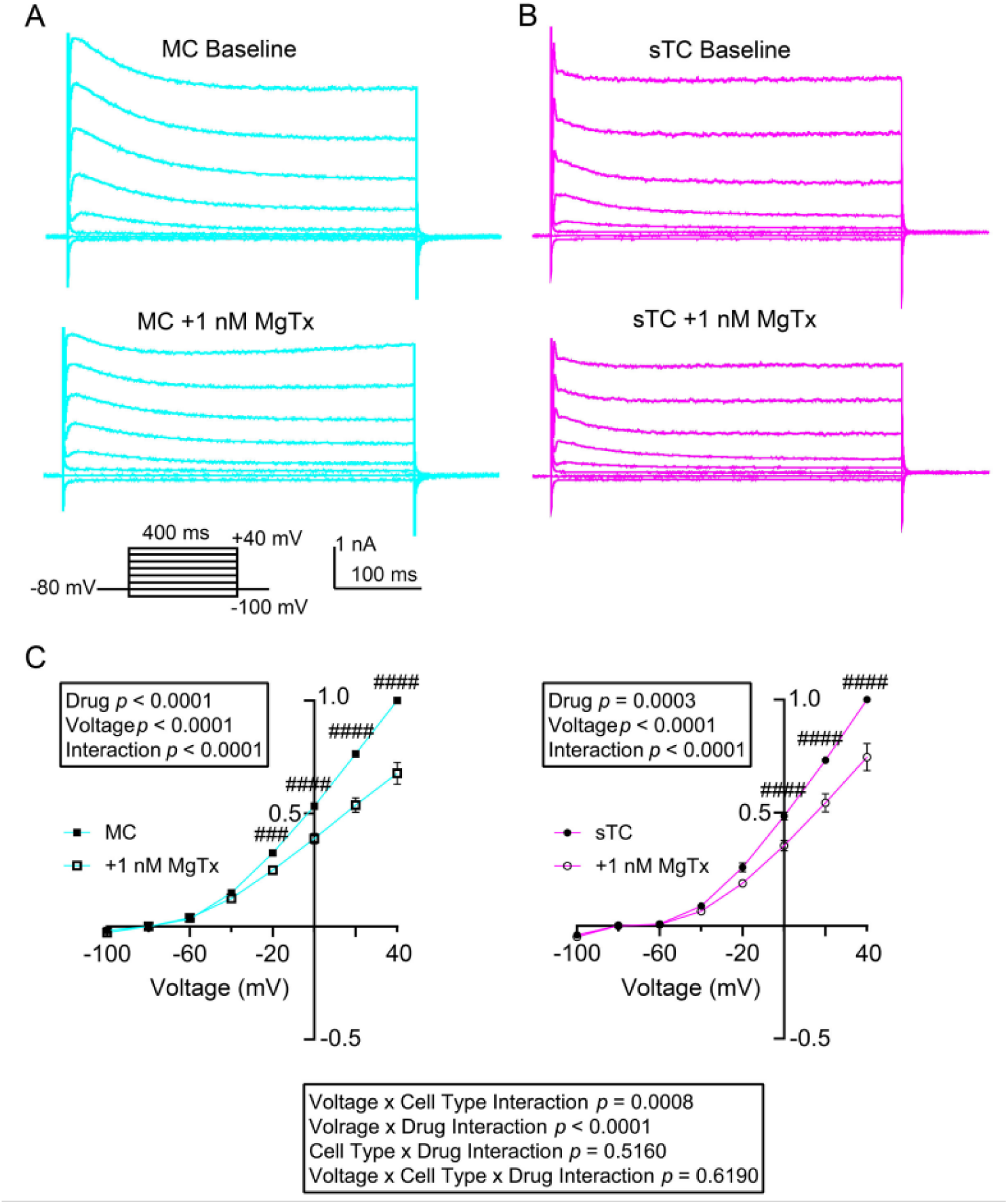
Sensitivity of mitral and superficial tufted cells to a selective blocker of Kv channels. ***A***, Family of voltage-activated currents recorded as in Figure 3 prior (***Top*,** Baseline) and following (***Bottom*,** +1 nM MgTx) 15-min bath application of margatoxin (MgTx) in a representative ***A,*** mitral cell (MC) and ***B,*** superficial tufted cell (sTC). ***C,*** Normalized IV plots prior (solid symbols) and following (open symbols) bath application of MgTx in a population of MCs (***Left,*** 2-w mixed RM ANOVA, drug F (1, 9) = 43.62, \*\*\*\**p* < 0.0001, with a Sidak’s post hoc test, ^###^*p* = 0.0008, ^####^*p* < 0.0001; voltage x drug interaction F (7, 63) = 43.86, \*\*\*\**p* < 0.0001) and a population of sTCs (***Right,*** 2-w mixed RM ANOVA, drug \*\*\**p* = 0.0003, Sidak’s post-hoc test, ^####^*p* < 0.0001; voltage x drug interaction \*\*\*\**p* < 0.0001). ***C, Bottom –*** 3-way mixed RM ANOVA result comparing the two IV plots in ***C, Left-Right*** (3-w mixed RM ANOVA, voltage x cell type interaction F (7, 123) = 3.879, \*\*\**p* = 0.0008; voltage x drug interaction F (7, 123) = 57.42, \*\*\*\**p* < 0.0001; cell type x drug interaction F (1, 123) = 0.4244, *p* = 0.5160; voltage x cell type x drug interaction F (7, 123) = 0.7633, *p* = 0.6190).

### Superficial tufted cells are more excitable than mitral cells

We next wanted to compare evoked AP firing frequency between MCs and sTCs as a metric for cellular excitability. After obtaining the whole-cell configuration, cells were current-clamped using a V_h_ of -60 mV (near RMP) for comparison of excitability across cell types. Each MC or sTC was stimulated with a family of injected current steps that increased in 25 pA increments from -50 pA to 175 pA (**Fig. 4A**) to determine the threshold for first firing. On average, the current needed to evoke APs (rheobase) was lower in sTCs than in MCs (**Fig. 4B**, MCs = 69.3 ± 44.9 pA (22) vs. sTCs = 34.2 ± 24.1 pA (30), Mann-Whitney U, \*\**p* = 0.0021). Both cell types increased AP firing frequency in response to current step, but sTCs had a significantly greater frequency than that of MCs (**Fig. 4C**). Here, we applied a 2-w mixed RM ANOVA using cell type and current as factors. As anticipated, there was a main effect for injected current (F(7, 263) = 71.39, \*\*\*\**p* < 0.0001). There was also a main effect for cell type (F (1, 51) = 11.92, \*\**p* = 0.0011) and a current x cell type interaction (F (7, 263) = 3.656, \*\*\**p* = 0.0009). A Sidak’s post-hoc test indicated a significant difference in current dependence at the current injection window between 25 and 75 pA (25 pA - ^$$^*p* = 0.0077, 50 pA - ^$$^*p* = 0.0017, and 75 pA - ^$$$^*p* = 0.0001). The increased AP firing frequency in response to injected current was likely attributed to a change in interspike interval (ISI). As shown in **Fig. 4D**, the ISI was shorter in sTCs compared to that of MCs. A 2-w mixed RM ANOVA using cell type and injected current as factors demonstrated a main (negative) effect for injected current (F (1.643, 38.07) = 33.74). There was also a main effect for cell type (F (1, 49) = 37.53, **** *p* < 0.0001) and a current x cell type interaction (F (6, 139) = 5.873, \*\*\*\**p* < 0.0001). A Sidak’s post-hoc test indicated a significant difference between cell types at the current injection window between 100 and 150 pA (100 pA - ^#^*p =*0.0101, 125 pA - ^##^*p =*0.0022, and 150 pA - ^#^*p =*0.0294).

**Fig. 4.**
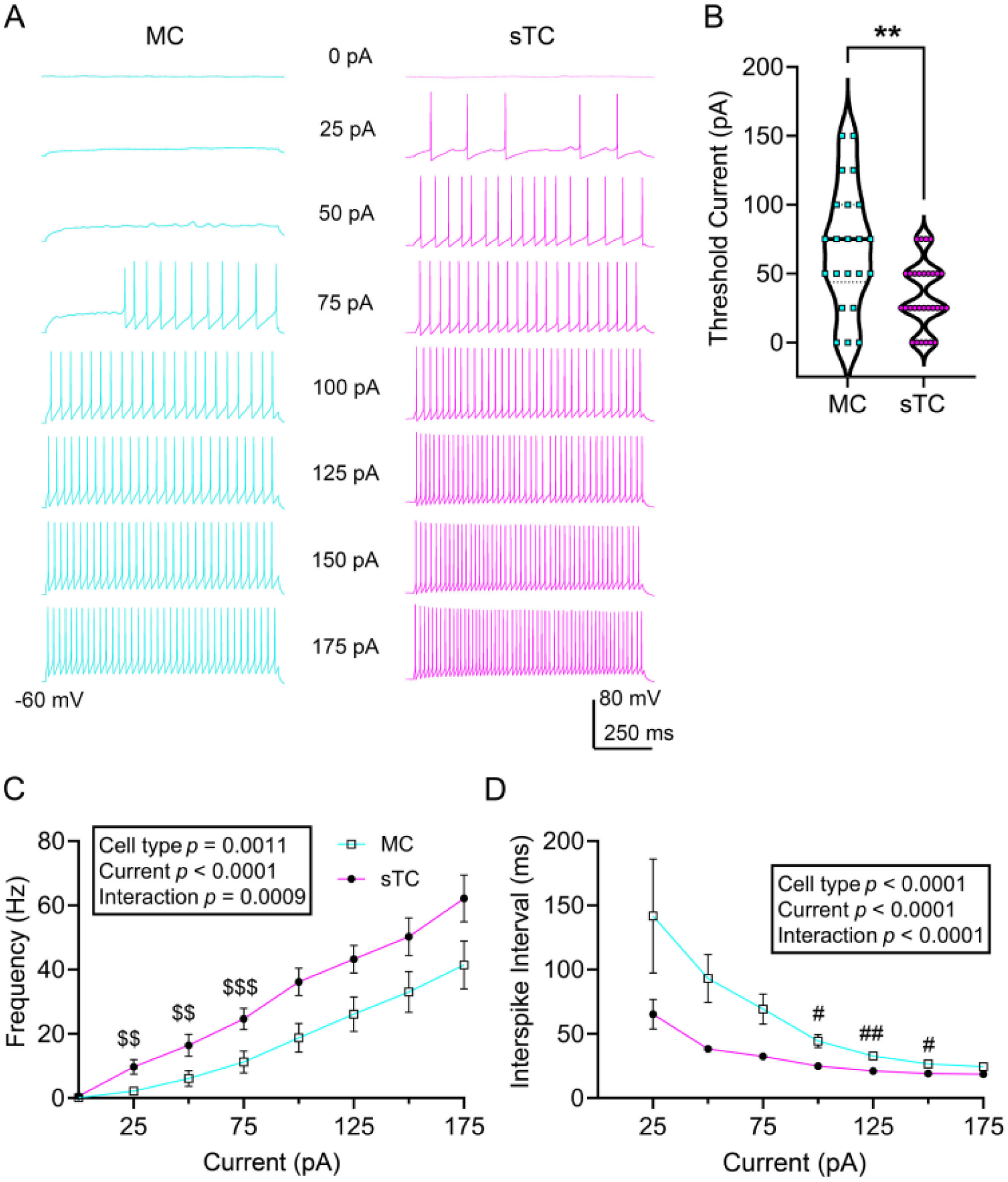
Spike firing properties compared between mitral cells (MC) and superficial tufted cells (sTC). ***A*,** Representative action potential trains evoked by various current steps as indicated, recorded from a MC (***Left***) vs. sTC (***Right***). ***B,*** Violin plot comparing the current needed to evoke action potentials (rheobase), Mann-Whitney U, \*\**p* = 0.0021. Line plots for ***C,*** mean spike frequency across current step and ***D,*** mean interspike interval (ISI). Line plot data were analyzed with a ***(C)*** 2-w mixed RM ANOVA, cell type \*\**p* = 0.0011; current x cell type interaction \*\*\**p* = 0.0009, Sidak’s post-hoc test, ^$$^*p* < 0.01, ^$$$^*p* < 0.001, ***(D)*** 2-w mixed RM ANOVA, cell type \*\*\*\**p* < 0.0001; current x cell type interaction \*\*\*\**p* < 0.0001, Sidak’s post-hoc test, ^#^*p* < 0.05, ^##^*p* < 0.01.

Because potassium channel currents can greatly influence the characteristics of APs (**Fig. 2-4**), we wanted to determine any differences in AP shape or kinetics that could accompany these findings. Many AP parameters require a clearly defined baseline in order to be assessed accurately, and the parameters are dependent on current injection magnitude. The metrics of ‘maximum decay slope’ and ‘time to achieve maximum decay slope’ are not dependent upon correct identification of the recording baseline, and we therefore examined these metrics simultaneously across all current injection steps. sTCs achieved steeper maximum decay slopes than MCs, which also differed in a current-dependent manner (**Fig. 5A, C**). We applied a 2-w mixed RM ANOVA using cell type and current step as factors. There was a main effect of cell type (F (1, 50) = 5.914, \**p* = 0.0186) and a current x cell type interaction (F (6, 127) = 5.562, \*\*\*\**p* < 0.0001); a Sidak’s post-hoc test indicated that cell-type significant differences occurred at current steps 50 pA - *p*^###^ = 0.0006, 75 pA - ^####^*p* < 0.0001, and 100 pA - ^##^*p* = 0.0074. APs recorded in sTCs also exhibited faster times to achieve maximum decay slope than those in MCs, independent of current injection step (**Fig. 5B, C**). We applied a 2-w mixed RM ANOVA using cell type and current step as factors. There was no effect of current step (F (0.1425, 3.016) = 9.085, *p* = 0.0524), but there was a main effect of cell type F (1, 50) = 4.088, \**p* = 0.0486. A Sidak’s post-hoc test, demonstrated that sTCs were faster at all current injections at and above 75 pA (75 pA - ^###^*p* = 0.0002, 100 pA - ^##^*p* = 0.0031, 125 pA - ^#^*p* = 0.0300, 150 pA - ^#^*p =* 0.0274).

**Fig. 5.**
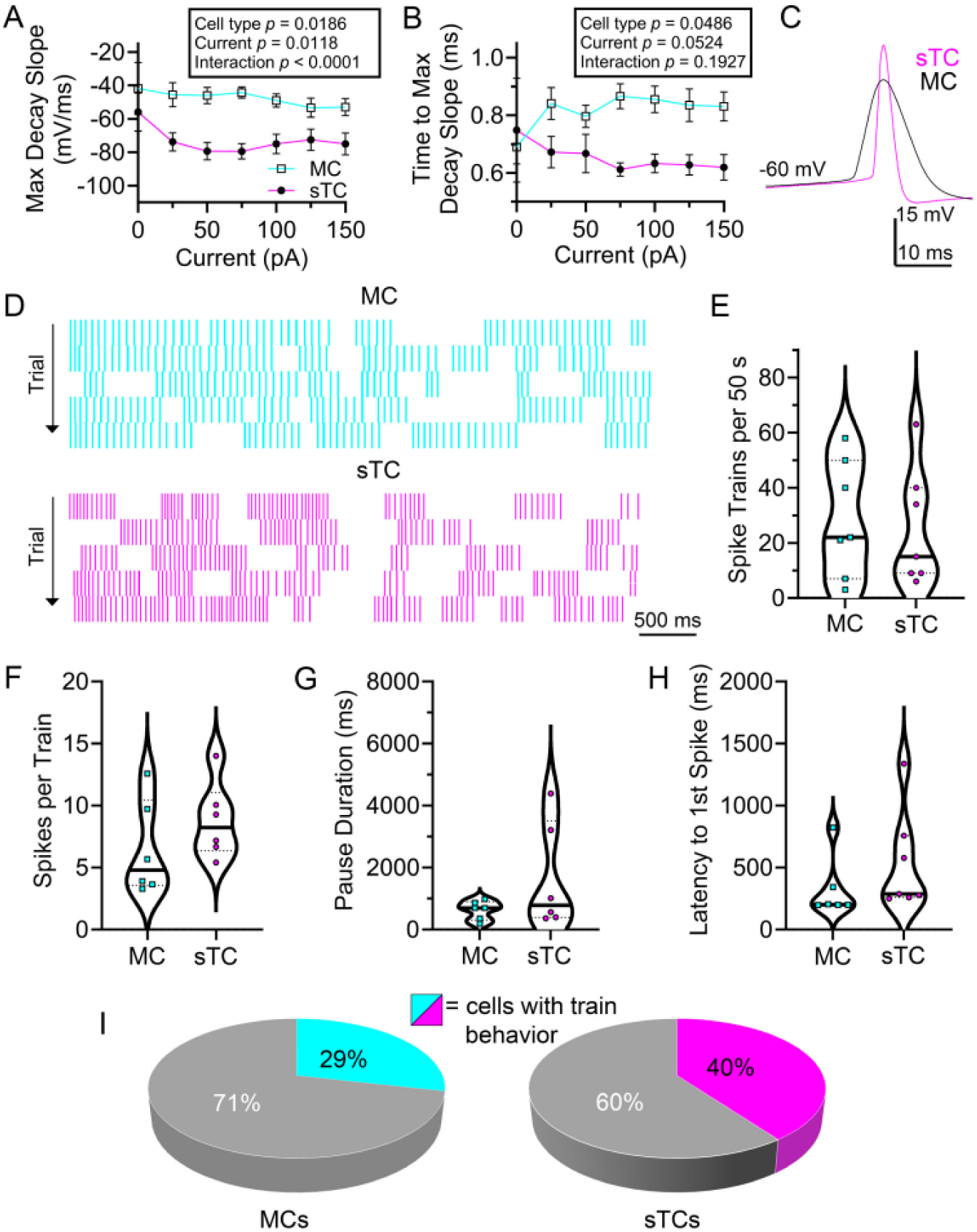
Action potential shape and spike train activity compared between mitral cells (MC) and superficial tufted cells (sTC). Line plot for ***A,*** maximum decay slope (max decay slope) and ***B,*** time to achieve maximum decay slope from peak magnitude (time to max decay slope) across current step. Line plot data were analyzed with **(A),** 2-w mixed RM ANOVA, cell type *p = 0.0186, Sidak’s post-hoc test, ##p < 0.01, ###p < 0.001; current x cell type interaction ****p < 0.0001, **(B),** 2-w mixed RM ANOVA, cell type *p = 0.0486, Sidak’s post-hoc test, #p < 0.05, ##p < 0.01, ###p < 0.001. ***C,*** Representative pair of action potentials recorded in response to a 75 pA current injection held near rest (V_h_ = -60 mV). ***D,*** Representative raster plot for a cell stimulated with a 50-pA current injection using a P_d_ = 5 s for each cell type, respectively. Successive trials are stacked vertically below. Violin plots computed across a 50 s recording period (10, 5s sweeps) compare ***E,*** mean number of bursts, ***F,*** mean spikes per train, ***G,*** mean pause duration between trains, and ***E,*** mean latency to first spike. ***E-F,*** were analyzed using a Student’s *t*-test, ***G-H*** were analyzed using Mann-Whitney U; *p* > 0.05. ***I,*** Relative proportion of cells eliciting spike train behavior among sampled MCs (***Left***) and sTCs (***Right***), respectively (Chi-squared test, p = .40061, X^2^ = 0.7065, n = 51).

Due to differences in excitability between MCs and sTCs (**Fig. 4**), the current injection step of 75 pA had the largest sample size for the comparison of AP characteristics that are dependent upon verifiable baseline. Overall, we found that the APs of sTCs have greater peak magnitude, greater antipeak (after-hyperpolarization) amplitude, faster rise kinetics, and faster decay kinetics than MCs (**Table 1**). Mean values, sample sizes, *p* values, and statistical tests are reported within each column.

**Table 1.**
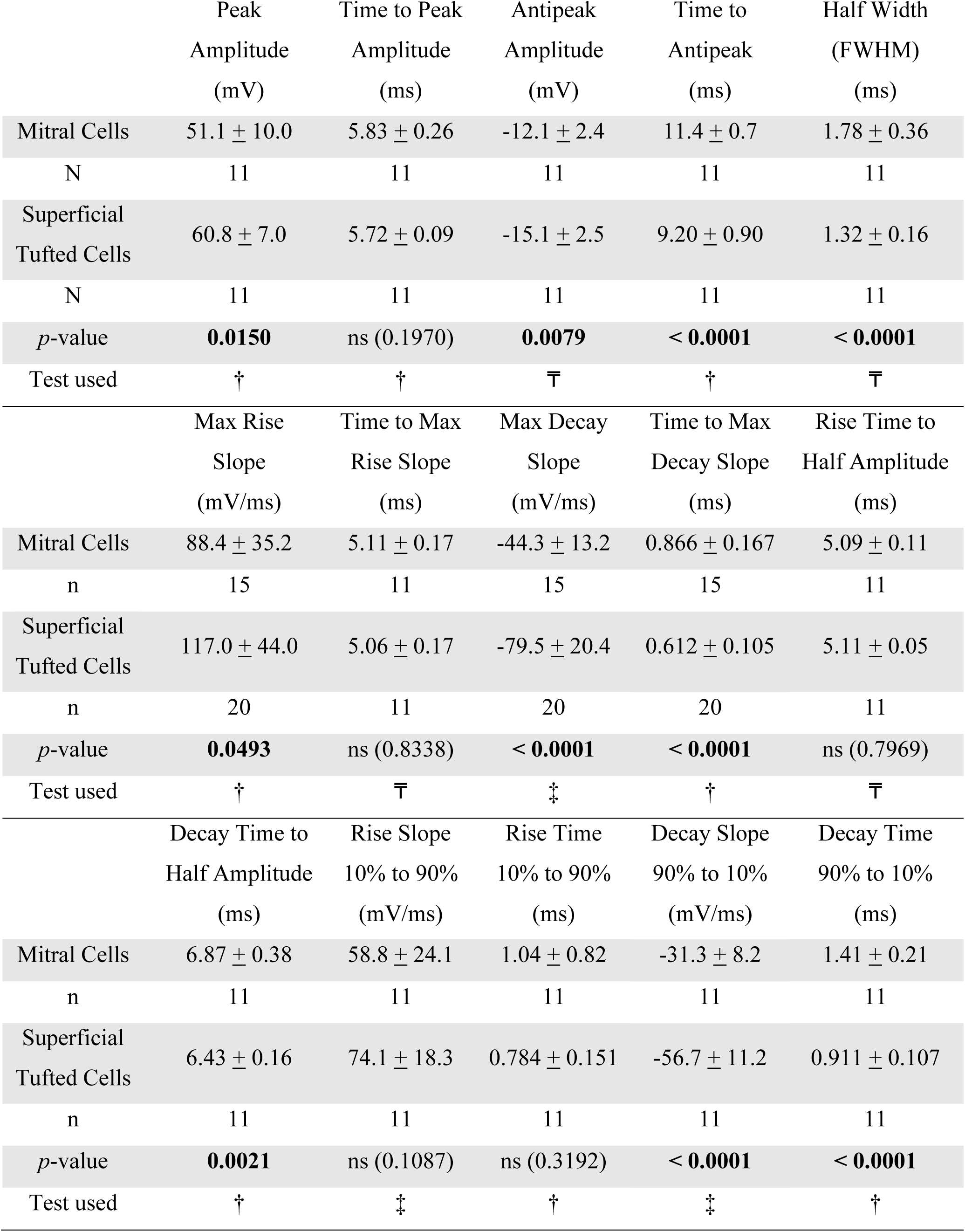
Action potential shape compared between mitral cells (MCs) vs. superficial tufted cells (sTCs). Recordings were made while holding cells near rest (V_h_ = -60 mV) and injecting 75 pA of current. Values are mean ± standard deviation. ns = not significant, †= Student’s *t*-test, ‡= *t*-test with Welch’s correction, ₸= Mann-Whitney U.

### Superficial tufted cells do not show a higher propensity to bursting activity than mitral cells

Previous reports demonstrate that MCs are less likely to exhibit trains of AP activity than TCs in general (Liu and Shipley, 2008; Burton and Urban, 2014; Jones *et al*., 2020). As such, we wanted to determine if this was also true of sTCs by applying further refined spike analysis. After determining the supra-threshold current injection step, we injected cells at that threshold using a P_d_ of 5000 ms and a T_s_ = 30 s for 10 total sweeps (**Fig. 5D**). We defined a “train” as having 3 or more APs within a 100 ms period (Balu et al., 2004), and only directly compared cells receiving identical current injection magnitude. sTCs did not exhibit more AP trains within their respective recording periods (**Fig. 5E**, MCs = 28.7 ± 21.1 (7) AP trains vs. sTCs = 25.1 ± 21.3 AP trains (7), Student’s *t*-test, *p* = 0.7582), nor did the trains of sTCs contain more AP spikes (**Fig. 5F**, MCs = 10.8 ± 11.9 spikes/train (6) vs. sTCs = 11.3 ± 7.2 spikes/train (6), Student’s *t*-test, *p* = 0.2796). sTCs did not exhibit differing pause duration between trains (**Fig. 5G**, MCs = 950 ± 890 ms (6) vs. sTCs = 1660 ± 1720 ms (6), Mann-Whitney U, *p* = 0.3939), nor a differing latency period to first spike (**Fig. 5H**, MCs = 327 ± 249 ms (6) vs. sTCs = 535 ± 404 ms (7), Mann-Whitney U, *p* = 0.1375). To assess whether sTCs showed a higher propensity to be a ‘bursting cell’ (as opposed to a regularly-firing cell), we assessed the firing activity from all current-clamp recordings to determine the categorical type of each MC and sTC. A cell was categorized as a “bursting cell” if any of its sweeps contained at least two bursts separated by a pause duration of > 100 ms. sTCs did not show a statistically different preference toward bursting or regularly firing than that of MCs (**Fig. 5I**, MCs = 28.6% (21), sTCs = 40% (30), Chi-squared test, *p* = 0.40061, X^2^ = 0.7065).

### Co-localization of Kv1.3 channel and the insulin receptor in superficial tufted cells

MCs are known be modulated by insulin, and Kv1.3 is necessary for this modulation (Fadool and Levitan, 1998; Fadool et al., 2000, 2011; Tucker et al., 2010; Kuczewski et al., 2014). We found sTCs to be modulated by margatoxin (**Fig. 3**), suggesting expression of Kv1.3. We therefore performed immunocytochemistry to determine co-expression of Kv1.3 and the insulin receptor (IR) in sTCs, similar to what we have observed for MCs (Marks and Fadool, 2007; Colley et al., 2009; Marks et al., 2009; Thiebaud et al., 2016). Using the GFP mice (Tbx21/Cas9/GFP), we observed the GFP-label localized to the M/TCs neurolamina (**Fig. 6A**, MCL and EPL, blue arrow MCs and orange arrow sTCs) as previously reported for this reporter line (Kolling et al., 2022). A majority of these neurons demonstrated expression of Kv1.3 channel and IR kinase (**Fig. 6B, 6C**) that was co-localilzed (**Fig. 6D**, white arrowhead) and overlayed with the GFP expression (merge, **Fig. 6E**). Correlated, higher magnification images were acquired to resolve the expression patterns and co-expression at the single neuron level (mitral cells **Fig. 6A1-E1**, superficial tufted cells **Fig. 6A2-6E2**).

**Fig. 6.**
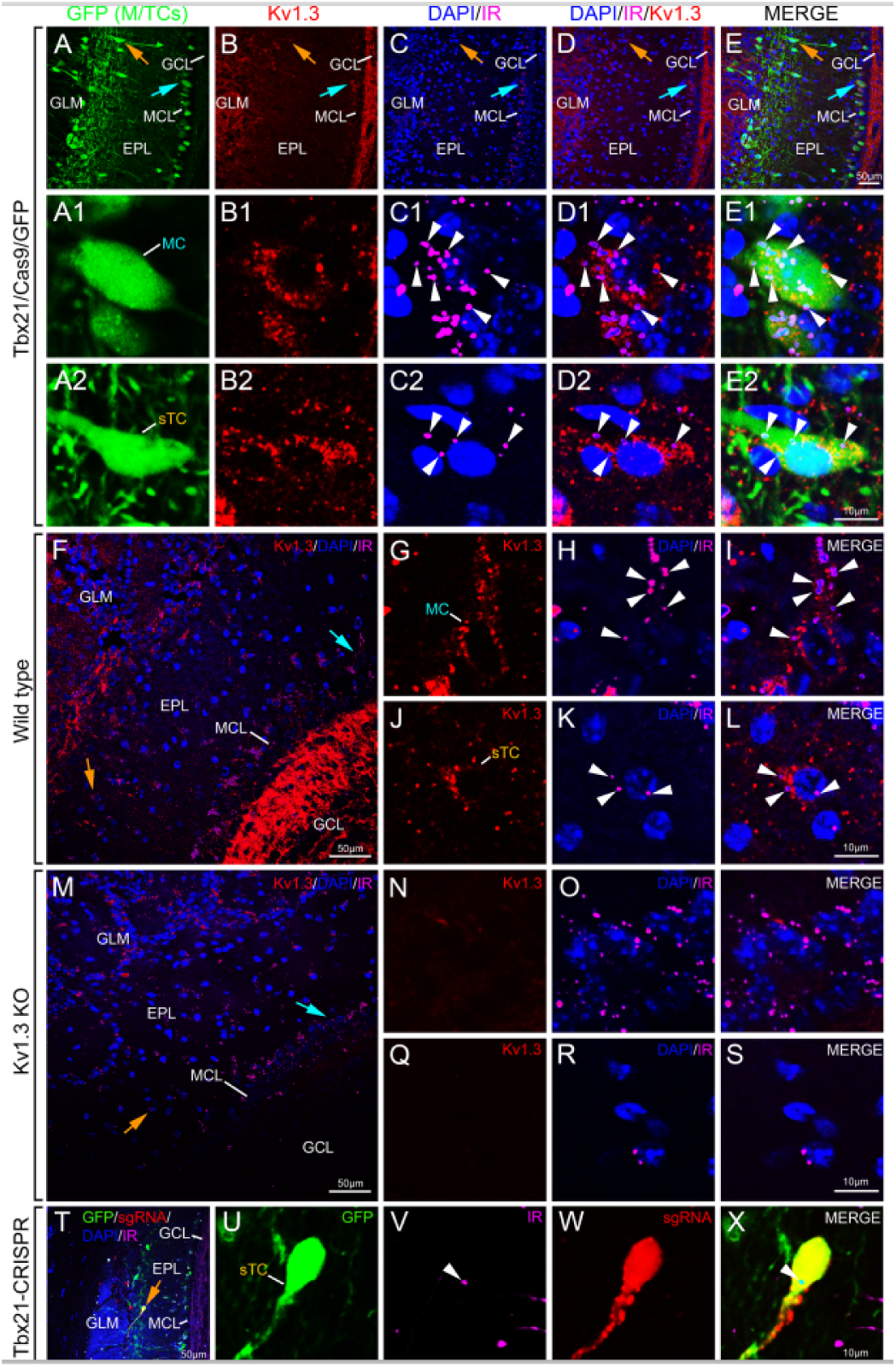
Colocalization of the Kv1.3 channel (Kv1.3) and the insulin receptor (IR) in superficial tufted cells (sTCs) and mitral cells (MCs) using double-color immunofluorescence confocal microscopy. Photomicrographs demonstrating Kv1.3- and IR-immunoreactivity in GFP mice (Tbx21/Cas9/GFP mice). ***A-E,*** Representative olfactory bulb slice at lower magnification indicating ***A,*** endogenous EGFP (GFP) selective in Tbx21 x Cas9 lines (green) ***B,*** Kv1.3 (red), ***C,*** IR (pink) and DAPI (blue) nuclear stain, and ***D,*** co-localization of Kv1.3 + IR, ***E,*** merge to GFP signal. ***A1-E1,*** Representative MC and ***A2-E2,*** representative sTC at higher magnification. GLM = glomerular layer, EPL= external plexiform layer, MCL = mitral cell layer, GCL: granule cell layer. Light blue arrow = MCL, orange arrow = sTCs near adjacent EPL, white arrowhead = Kv1.3 + IR co-labeled neuron. Representative photomicrographs demonstrating Kv1.3- and IR-immunoreactivity in ***F,*** wildtype and ***M,*** Kv1.3-/- mice. ***G-I,*** Representative MC and ***J-L,*** representative sTC at higher magnification for wildtype mice in comparison with a ***N-P,*** MC and a ***Q-S,*** sTC for Kv1.3-/- mice. Abbreviations and notations the same as (**A-E2**). ***T-Y,*** Representative photomicrographs demonstrating IR-immunoreactivity following CRISPR editing of Kv1.3 channel (Tbx21-CRISPR). ***U,*** Endogenous EGFP (GFP) selective in Tbx21 x Cas9 lines, ***V,*** IR, ***W,*** sgRNA, ***X,*** higher magnification of a sTC, merge to GFP signal and ***T,*** associated low power of identical field of view. Abbreviations and notations the same as (**A-E2**).

Using Kv1.3-/- mice as a negative control for channel expression and co-localization with IR kinase, we observed that the prominent detection of individual protein label and co-expression patterns observed in the MCL, EPL, and GCL of wildtype mice (**Fig. 6F**), were absent in Kv1.3-/- mice (**Fig. 6M**). Higher magnification images showing labeling in wildtype mice (mitral cells **Fig. 6G-I**; superficial tufted cells **Fig. 6J-L**) vs. that of Kv1.3-/- mice (mitral cells **Fig. 6N-P**; superficial tufted cells **Fig. 6Q-S**) further resolved the lack of signal for the Kv1.3 channel. These data indicated that loss of Kv1.3 channel did not grossly affect expression patterns of IR kinase. To further confirm the retention of IR kinase expression patterns in sTC following loss of Kv1.3 expression, we examined GFP mice following CRISPR-mediated deletion of Kv1.3 channel in M/TCs (**Fig. 6T-6X**). Indeed, sTCs that had been CRISPR-edited – expressing Cas9 (**Fig. 6U**) and taking up sgRNA (**Fig. 6W),** continued to express IR kinase (**Fig. 6V**) in the absence of Kv1.3 channel (merge low power **Fig. 6T**, high magnification **Fig. 6X**).

We next performed multiplex fluorescent *in situ* hybridization (RNAscope) to determine co-expression of Kv1.3 and the insulin receptor (IR) in sTCs as a secondary confirmation with resolution at the molecule level to include quantification. Using a transgenic GFP label for M/TCs under control of the *Tbx21* promoter, we performed RNAscope targeting *Kv1.3*, *GFP*, and *Insr* on six coronal sections sampled 512 µm apart along the antero-posterior axis (AP axis), in both male and female mice. We observed co-localization of the channel and the receptor markedly distributed with the expression of the *Tbx21* promoter by imaging at low power across the entire olfactory bulb (**Fig 7A-D**). We then moved to higher magnification in the EPL to visualize the sTC specifically, and the colocalization of the channel and receptor with the *Tbx21* reporter expression (**Fig. 7E-H**). A 2-way comparison revealed a main effect of AP axis on Tbx21+ cell count (**Fig. 7I**, \**p* = 0.0118) with no interactive effect of axis on sex or laminar layer (*p* = 0.3020; M MCL, M EPL, M GLM, F MCL, F EPL, F GLM as columns variables; M = male, F = female). We then examined the percentage of Tbx21+ cells co-labeled with both *Kv1.3* and *Insr* along the antero-posterior axis, and found no main effect of AP axis (**Fig. 7**, *p* = 0.5736) or interactive effects on sex or laminar layer (*p* = 0.7359).

**Fig. 7.**
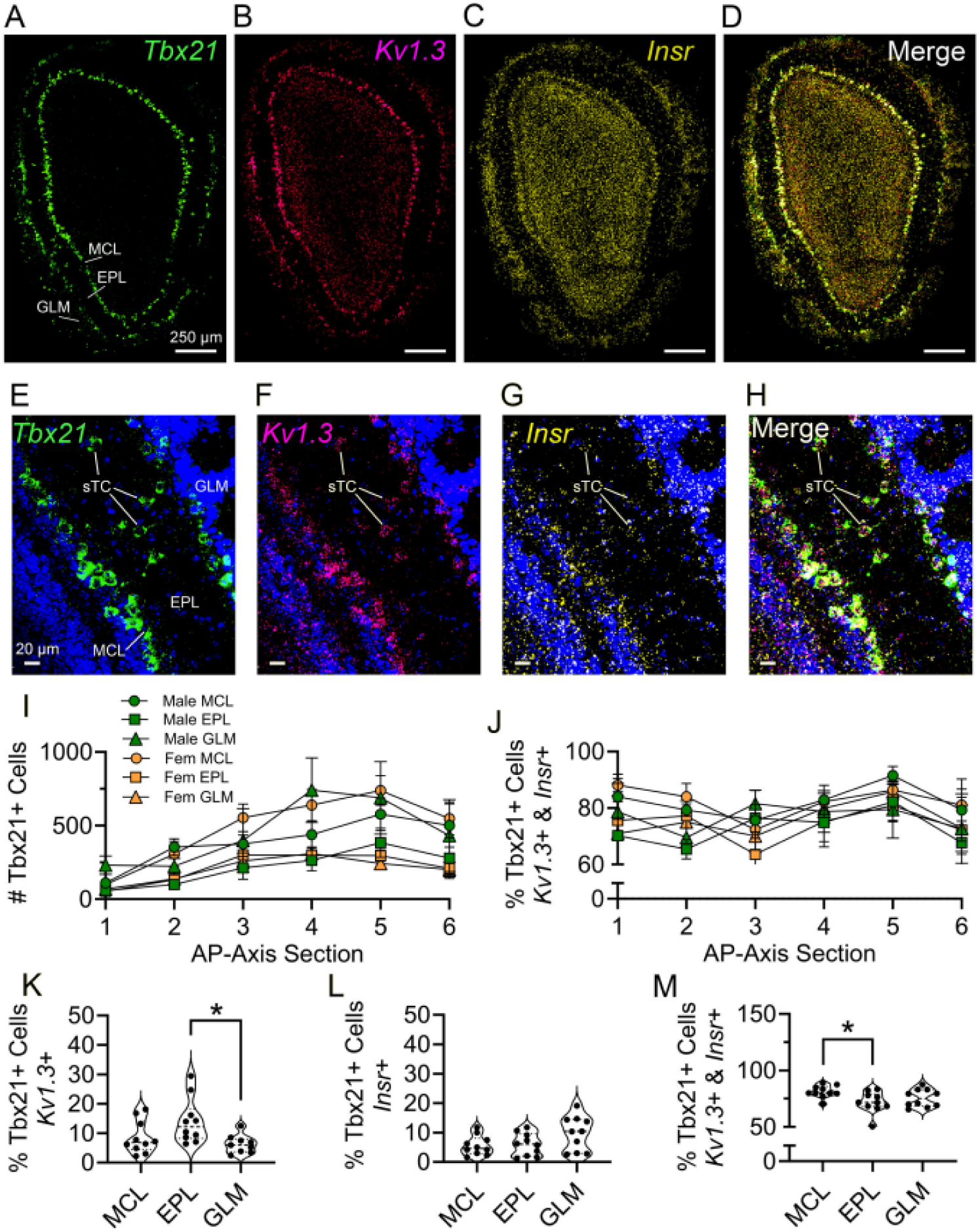
Colocalization of MCs/sTCs with the Kv1.3 channel (*Kv1.3*) and insulin receptor (*Insr*) using RNAscope. Photomicrographs demonstrating neurolaminar distribution of Tbx21 reporter expression (green, ***A***), *Kv1.3* expression (magenta, ***B***), and *Insr* expression (yellow, ***C***), separately and merged (white, ***D***). ***E–H***: Same as ***A–D***, but to demonstrate localization to sTCs with the inclusion of a DAPI nuclear stain. ***I:*** Coronal sections were sampled along the anter-posterior axis (AP axis) at ∼512 µm intervals and assessed for total number of M/sTCs (Tbx21+ cells) and for percentage of M/sTCs expressing both *Kv1.3* and *Insr* (***J***). Each data point represents the average from 5 separate mice, separated by laminar layer and sex. ***K–M:*** Percentage of M/TCs (Tbx21+ cells) expressing *Kv1.3* alone (***K***), *Insr* alone (***L***) or both (***M***). Statistical comparisons for ***K–M*** are Kruskal-Wallis (ANOVA) with Dunn’s multiple comparisons test. Each data point represents the average of all six antero- posterior sections from each mouse, one data point per mouse per neurolaminar layer, respectively. \**p* < 0.05, GLM = glomerular layer, EPL = external plexiform layer, GCL = granule cell layer.

Without regard for antero-posterior position or sex (since these were not found to be factors for co-labeling), we assessed the percentage of Tbx21+ cells that express both *Kv1.3* and *Insr*. We find triple-labeling of 82.1 ± 4.3 % in the MCL, 73.4 ± 8.2 % in the EPL, and 76.2 ± 6.8 % in the GLM. ANOVA comparisons across neurolaminae revealed a subtle difference in the percentage of all Tbx21+ cells expressing *Kv1.3* alone (**Fig. 7K**, \**p* = 0.0159) and expressing both *Kv1.3* and *Insr* (**Fig. 7M**, \**p* = 0.0373).

### Superficial tufted cells are modulated by insulin via the Kv1.3 ion channel

To further explore the complementary physiological role of sTCs, we wanted to determine if their excitability could be modulated by insulin—knowing that MCs are modulated by 0.1 µm insulin (Fadool *et al*., 2000, 2011). After sampling for supra-threshold current, synaptic blockers were added to the bath solution. Evoked APs were established using the threshold current with a P_d_ = 5000 ms and T_s_ = 30 s for a 5-min paradigm (10 recordings). A representative set of baseline recordings and an associated raster plot is shown in **Fig. 8A**, prior (left) and subsequent (right) to bath application of insulin for 15 min.

**Fig. 8.**
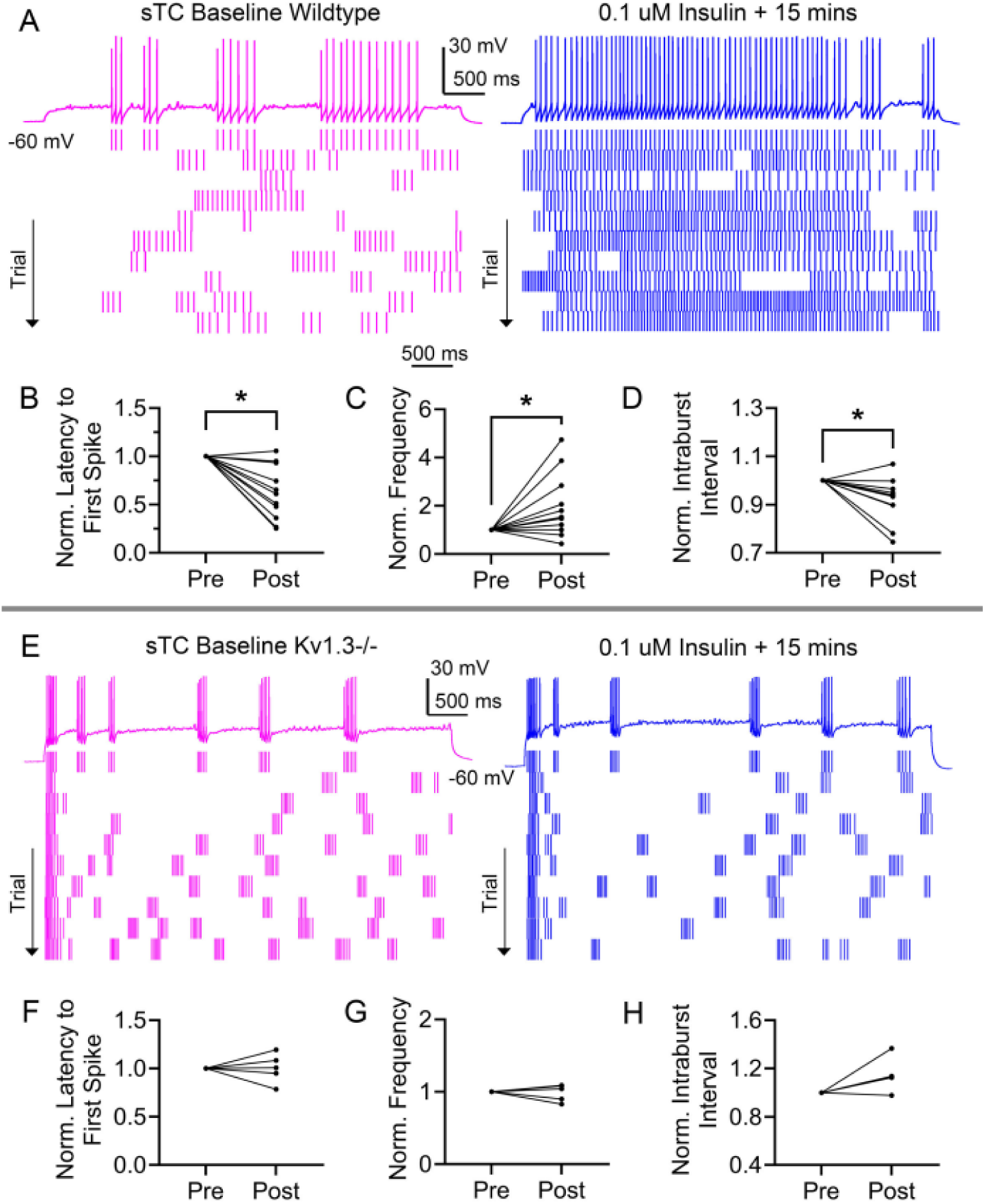
Bath application of insulin increases spike firing frequency, and shortens latency to first spike and intraburst interval in superficial tufted cells (sTCs) in wildtype mice (***A-D***), but fails to do so in superficial tufted cells in Kv1.3-/- mice (***E-H***). ***A,*** Representative spike trains (***Top***) that were elicited in response to 25 pA current injection while holding the cell near rest (V_h_ = -60 mV) prior (***Left,*** Baseline) and subsequent (***Right,*** Insulin) to hormone stimulation using P_d_ = 5 s. (***Bottom***) Representative associated raster plot. Line graph comparing ***B,*** latency to first spike, ***C,*** spike firing frequency, and ***D,*** intraburst interval, prior (pre) and following (post) insulin stimulation in the same cell. Data were analyzed with a paired *t*-test using non-normalized data ***(B)*** \**p* = 0.0132, ***(C)*** \**p* = 0.0381, and ***(D)*** \**p* = 0.0136. ***E,*** Representative spike trains (***Top***) and associated raster plot (***Bottom***) recorded as in *Panel A,* but in Kv1.3-/- mice. Line graph comparing ***F,*** latency to first spike, ***G,*** spike firing frequency, and ***H,*** intraburst interval, prior (pre) and following (post) insulin stimulation in the same cell. Data were analyzed with a paired *t*-test using non-normalized data, *p* > 0.05.

Insulin significantly reduced the average latency to first spike by ∼38% (**Fig. 8B**, sTC pre = 1220 ± 723 ms (11) vs. sTC post = 649 ± 403 ms (11), paired *t*-test, \**p* = 0.0132). On average, insulin also increased the AP frequency by ∼98% (**Fig. 8C**, sTC pre = 4.76 ± 1.76 Hz (11) vs. sTC post = 8.49 ± 4.98 Hz (11), paired *t*-test, \**p* = 0.0381), and decreased the average intraburst interval by ∼8% (i.e., the pause between APs within a particular burst) (**Fig. 8D**, sTC pre = 65.1 ± 13.6 ms (11) vs. sTC post = 60.3 ± 14.9 ms (11), paired *t*-test, \**p* = 0.0136). All statistical tests were performed on non-normalized data, however, plots were generated as normalized paired values to visualize individual cell changes across the sampled cells.

The modulation of MCs by insulin has been shown to depend on the Kv1.3 ion channel using either pharmacological block or targeted gene-deletion (Fadool *et al*., 2000, 2011; Colley *et al*., 2004). Since previous reports demonstrate Kv1.3 expression in sTCs (Kues and Wunder, 1992; Fadool *et al*., 2000; Colley *et al*., 2007), we wanted to determine if insulin modulation of sTCs was also Kv1.3-dependent. Using transgenic mice lacking the *Kv1.3* gene (Kv1.3-/-, see methods) we first compared the AP bursting characteristics between the sTCs of GFP mice (Kv1.3 +/+), and the sTCs of Kv1.3 -/- mice. In accordance with the findings in Kv1.3 - /- MCs (Fadool *et al*., 2004, 2011; Tucker *et al*., 2010; Thiebaud *et al*., 2016), we found that Kv1.3 -/- sTCs tend to burst more often, and have shorter bursts with faster intraburst frequency. Kv1.3 -/- sTCs also have shorter latency to first spike and faster overall evoked AP frequency than Kv1.3 +/+ sTCs (**Table 2**). Mean values, sample sizes, *p* values, and statistical tests are reported within each column.

**Table 2.**
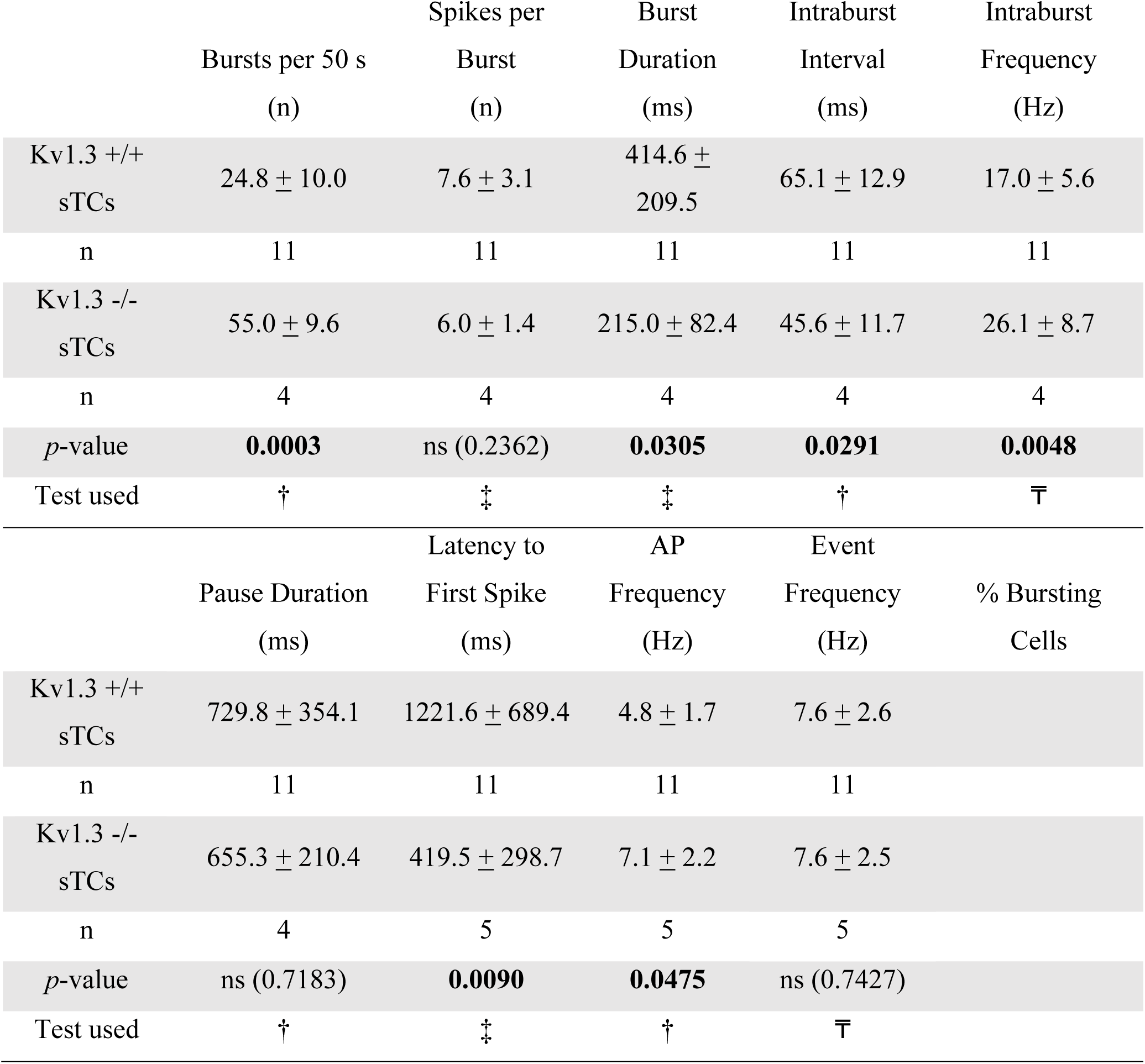
Spike bursting properties in superficial tufted cells (sTCs) compared between wildtype (WT) and Kv1.3 -/- mice. Recordings were made while holding cells near rest (V_h_ = -60 mV) and injecting 75 pA of current. Data were computed across a 50 s recording period (10, 5s sweeps). Values are mean ± standard deviation. ns = not significant, †= Student’s *t*-test, ‡= *t*-test with Welch’s correction, ₸= Mann-Whitney U

|  | Bursts per 50 s<br>(n) | Spikes per<br>Burst<br>(n) | Burst<br>Duration<br>(ms) | Intraburst<br>Interval<br>(ms) | Intraburst<br>Frequency<br>(Hz) |
| --- | --- | --- | --- | --- | --- |
| Kv1.3 +/+<br>sTCs | 24.8 ± 10.0 | 7.6 ± 3.1 | 414.6 ±<br>209.5 | 65.1 ± 12.9 | 17.0 ± 5.6 |
| n | 11 | 11 | 11 | 11 | 11 |
| Kv1.3 -/-<br>sTCs | 55.0 ± 9.6 | 6.0 ± 1.4 | 215.0 ± 82.4 | 45.6 ± 11.7 | 26.1 ± 8.7 |
| n | 4 | 4 | 4 | 4 | 4 |
| p-value | **0.0003** | ns (0.2362) | **0.0305** | **0.0291** | **0.0048** |
| Test used | † | ‡ | ‡ | † | ⎯ |
|  | Pause Duration<br>(ms) | Latency to<br>First Spike<br>(ms) | AP<br>Frequency<br>(Hz) | Event<br>Frequency<br>(Hz) | % Bursting<br>Cells |
| Kv1.3 +/+<br>sTCs | 729.8 ± 354.1 | 1221.6 ± 689.4 | 4.8 ± 1.7 | 7.6 ± 2.6 |  |
| n | 11 | 11 | 11 | 11 |  |
| Kv1.3 -/-<br>sTCs | 655.3 ± 210.4 | 419.5 ± 298.7 | 7.1 ± 2.2 | 7.6 ± 2.5 |  |
| n | 4 | 5 | 5 | 5 |  |
| p-value | ns (0.7183) | **0.0090** | **0.0475** | ns (0.7427) |  |
| Test used | † | ‡ | † | ⎯ |  |

Using Kv1.3 -/- mice, we next repeated the previous insulin experiment shown **Fig. 8A** representative set of baseline recordings and associated raster plot for sTCs evoked APs is shown in **Fig. 8E**, prior (left) and subsequent (right) to bath application of insulin for 15 min as recorded in the Kv1.3-/- mice. Unlike in recordings acquired from Kv1.3 +/+ mice, insulin did not alter latency to first spike (**Fig. 8F**, Kv1.3-/- sTC pre = 420 ± 334 ms (5) vs. Kv1.3-/- sTC post = 390 ± 232 ms (5), paired *t*-test, *p* = 0.5694). Insulin also did not affect AP firing frequency (**Fig. 8G**, Kv1.3-/- sTC pre = 7.09 ± 2.48 Hz (5) vs. Kv1.3-/- sTC post = 7.13 ± 2.93 Hz (5), paired *t*-test, *p* = 0.9300, n = 5), nor affect the intraburst interval (**Fig. 8H**, Kv1.3- /- sTC pre = 45.6 ± 13.6 ms (4) vs. Kv1.3-/- sTC post = 52.7 ± 17.8 ms (4), paired *t*-test, *p* = 0.1990, n = 4) in Kv1.3-/- mice. The same 5 cells were used for all pre/post comparisons, but one cell did not fire quickly enough for burst analysis (see methods).

## Discussion

We have demonstrated that the two major output neurons in the olfactory bulb have different biophysical properties, despite both neurons expressing IR kinase and exhibiting modulation by insulin. sTCs are more excitable, as evidenced by the lower threshold needed to evoke an AP, a decreased ISI, and a greater AP firing frequency particularly at lower current injection. We found that the APs of sTCs have greater peak amplitude, greater anti-peak amplitude, faster rise kinetics, and faster decay kinetics than those of MCs – while at the same time, the incidence of a train of AP activity was similar between the two output neurons. While there are differences in the voltage dependence of voltage-activated outward currents between MCs and sTCs, both are blocked by the Kv1.3 vestibule blocker, MgTx. We determined that voltage-activated outward currents in sTCs are modulated by insulin, and that the Kv1.3 ion channel is necessary for this modulation.

Burton and Urban (Burton and Urban, 2014) first compared mitral cell biophysical properties with that of what they called “tufted cells”, which were projection neurons whose cell bodies resided in the superficial 50% of the EPL. Prior to their publication, physiologists often clustered mitral and tufted cells together as a homogenous neuronal population (Fadool and Levitan, 1998; Tucker and Fadool, 2002; Youngstrom and Strowbridge, 2015; Liu et al., 2016) due to their similar function as output neurons. Using new nomenclature described by Jones et al. (Jones et al., 2020), we now know that TCs are comprised of a variety of sub-classes that are functionally distinct with differential sensitivity to inhibitory/excitatory circuits to putatively sharpen tuning specificity for odorants during sensory processing (Zhou et al., 2023). We also know from transcriptome and RNA sequencing studies that there are multiple subclasses of each of these projection neurons based upon molecular determinants (Zepelli et al., 2021). The tufted cells first measured by Burton and Urban (2014) likely consisted of a mixed population of sTCs and mTCs. Nonetheless, some of our data measured in strictly sTCs, align well with what they reported for “tufted cells” in general. Burton and Urban found that their TCs had a higher instantaneous frequency (peak instantaneous rate) that differed with current injection magnitude than that of MCs, and that the AP shape of their TCs exhibited a narrower half width (FWHM), a steeper decay slope, and faster time to anti-peak (after-hyperpolarization). Our comparative measurements were quite similar. Thus, we similarly report that sTC have greater intrinsic excitability than that of MCs. Unlike this earlier study, however, we found that the AP shape in sTCs compared with that of MCs, has a greater peak amplitude, steeper maximum rise slope, a greater anti-peak amplitude, and the minimum current necessary to evoke a first AP (threshold) is lower.

Burton and Urban further found that their TCs were more likely to elicit spike trains (stutter) compared with that of MCs, which they characterized based upon the increased co-efficient of variation of the ISI observed in the TCs. We did not observe a greater overall firing irregularity in our recorded sTCs, and in fact, we found the opposite - our recorded MCs had a greater ISI variance at lower levels of current injection than that of sTC. Also inconsistent with our findings; sTCs and MCs did not have significantly different AP train properties (spikes per train, pause duration, or latency to first spike), nor did sTCs have a statistically higher propensity to elicit a spike train. The likely inclusion of both sTCs and mTCs in their study could account for these differential findings. In fact, Antral et al. (2006) also reports in their study of eTC, that they observed two subclassifications, nonbursting (contained lateral dendrites) and bursting (lacked lateral dendrites), that were found to be both morphologically and functional diverse. We believe that we have complemented the findings of Burton and Urban, expanding upon their comparisons to more specifically characterize selectively the sTCs and not multiple subtypes clustered together.

Differences in ion channel expression could underlie the greater excitability of sTCs compared to that of MCs. We found voltage-activated currents in the window of -60 to -20 mV to be greater in MCs than those of sTCs. It is well known that *shaker* family members have a voltage-activation above -50 to -40 mV (Zagotta and Aldrich, 1990), therefore Kv1.2 and Kv1.3 likely are not highly responsible for the difference in voltage-activated currents between the two cell types. This is corroborated by the fact that both cell types exhibited a similar pharmacological response to MgTx and both are shown to be similarly modulated by bath application of insulin, previously demonstrated to be the result of Kv1.3 being used as a substrate for tyrosine phosphorylation in the OB in a use- or activity-dependent manner (Fadool et al., 2000, 2011). We also measured a select slowing of the deactivation kinetics in MCs at the sampled -40 mV hyperpolarizing step that might selectively lower MC excitability vs. that in sTCs. A comparison of MC activity in Kv1.3-/- mice vs. pharmacological block of the channel in the current study, may reveal differences in underlying ion channel expression between sTCs and MCs. The threshold for activation of voltage-activated outward currents in the MCs of Kv1.3-/- mice were previously found to be shifted +20 mV in the positive direction compared to wild-type MCs (Fadool et al., 2004), but these Kv1.3-/- MCs have an upregulation of a *slack* sodium-activated potassium (K_Na_) channel (Lu et al., 2010), an increase in calcium channel expression (Fadool et al., 2004), and may have increased chloride currents (Koni et al., 2003). Therefore, the threshold for activation of voltage-activated currents in sTCs and Kv1.3-/- MCs is both higher than that of wildtype MCs, but the precise nature of any differences in channel expression (i.e., perhaps sTCs express higher levels of K_Na_ channels than MCs) is not known. One study that has tried to address differential expression of voltage-gated K channels across MC and TCs, in general, is that of Zeppilli et al. (2021). These authors have demonstrated distinct RNA expression levels of Kcnd3, Kcng1, Kcnh5, Kcnq3, Kcnj2 and 6, and Hcn1 that may represent candidates for controlling the differential excitability of different MC and TC subtypes. This opens the door to a larger area of research that is beyond the scope of this report.

sTCs are modulated by the presence of insulin, and our data demonstrate that this modulation depends upon the Kv1.3 ion channel. Insulin’s primary role in the periphery is to mobilize glucose from the bloodstream following a meal; however, insulin is known to have many roles as a signaling molecule in the central nervous system (CNS). The levels of insulin in the CNS do not mirror the levels of insulin in the periphery, and the highest concentration of insulin receptors and binding affinity to insulin (outside the pancreas) is found in the OB (Baskin et al., 1983; Hill et al., 1986; Gupta et al., 1992; Schwartz et al., 1992; Myers Jr. and White, 1993; White, 1997; Wickelgren, 1998; Banks et al., 1999; Fadool et al., 2000; Aimé et al., 2012; Palouzier-Paulignan et al., 2012; Fadool and Kolling, 2020). MCs have an interesting co-localization of IR kinase and Kv1.3 with adaptor proteins that functions to cluster signaling transduction cascades in close proximity (Marks and Fadool, 2007; Marks et al., 2009). In the brain, insulin resistance is implicated in the progression of neurodegeneration and dementia (Kuljiš and Salković-Petrišić, 2011; Arnold et al., 2018; Ferreira et al., 2018; Hong et al., 2021; Femminella et al., 2021). Our report elevates the role of Kv1.3 as a target for the investigation of olfactory function, and, by extent, the complexities of insulin signaling in the OB. Though not annotated directly, previous histological reports suggested expression of Kv1.3 in sTCs. These include Figure 1E from Kues 1992 (Kues and Wunder, 1992), Figure 7 from Fadool 2000 (Fadool et al., 2000), and Figure 9 from Colley 2007 (Colley et al., 2007). At this time, we do not know of any reports that previously suggested expression of insulin receptors on sTCs. MCs are known to be modulated by insulin in a similar manner to our sTC findings (Fadool et al., 2000, 2011; Das et al., 2005; Tucker et al., 2010; Aimé et al., 2012; Kuczewski et al., 2014), and this modulation in MCs is known to occur via phosphorylation of Kv1.3 (Bowlby et al., 1997; Fadool and Levitan, 1998; Fadool et al., 2000, 2011).

Tbx21-cre lines have been previously utilized to selectively label M/TCs and delineate downstream olfactory pathways (Mitsui et al., 2011; Kolling et al., 2022). Herein, we utilized this line to parse out the TC subtypes based on their position in the neurolamina. Juxtaglomerular eTCs situated at the periphery of the glomeruli could be clearly distinguished from sTCs situated along the outer two thirds of the EPL (Jones et al., 2020). An autoradiographic insulin binding study (Hill et al., 1986) and *in situ* hybridization analyses (Marks et al., 1990) have confirmed the high density of insulin receptor expression across different layers of olfactory bulb. In agreement with our current immunocytochemical data, Aime et al. (2012) demonstrated the highest density of IR kinase specifically in the mitral cells of rats. We observed that the number of IR-immunoreactive puncta was higher in number and density in the MCL compared to that of the adjacent EPL and GLM. Interestingly, we could use the Tbx21-cre lines to visualize sTCs that expressed IR, and noted that the density of IR punta in sTCs was comparatively lower than that observed in MCs. Although we did not examine the mechanism by which insulin might regulate sTCs, it is predicted that Kv1.3 is a substrate for insulin-mediated phosphorylation (Fadool et al., 2000; 2011), which is in keeping with co-localization of the channel and the kinase in both MCs and sTCs. We note that despite reported signaling interaction between channel and kinase (Marks and Fadool, 2007; Marks et al., 2009), and the fact that modulation of spike firing frequency in sTCs was dependent upon the presence of Kv1.3 channel, IR expression was retained in both Kv1.3-/- and CRISPR-edited mice. While we do not know if the general insulin signaling pathway is perturbed in the absence of Kv1.3, expression of the receptor appears to be independent of the channel, and reciprocally, insulin modulation of the projection neurons is dependent upon the channel. In agreement with prior physiological findings (Tucker et al., 2010), Kv1.3-/- mice, lacking Kv1.3 immunoreactivity, also do not hamper the expression of IR across different OB layers.

While we know that IR kinase is also co-localized with glucagon-like 1 peptide receptors in MCs (Thiebaud et al., 2016) and have discovered an array of neuromodulators that alter Kv1.3 channel function and general excitability of these output neurons (Cook and Fadool, 2002; Tucker and Fadool, 2002; Colley et al., 2009; Fadool et al., 2011; Mast and Fadool, 2012; Tucker et al., 2013; Huang et al., 2017; Thiebaud et al., 2019), the possibility of neuromodulation of sTCs remains highly likely and greatly unexplored. Herein, we have distinguished these two output neurons in terms of biophysical function, but there may be a range of neuromodulators that can affect and fine tune odor information coding or regulation of metabolic state (Thiebaud et al., 2016, 2019; Kolling et al., 2022). Moreover, we have recently enhanced total excitability of the output neurons (MCs and all classes of TCs) using cell selective CRISPR genome editing of Kv1.3 to support a new function of the olfactory system – metabolic regulation and energy homeostasis (Kolling et al., 2022). Thus, there is much to discover concerning both the division of olfactory function between MCs and TCs (odor context, valence, vs. odor identity), as well as the combined parallel function of olfactory output neurons in regulating whole-body metabolism.

## MATERIALS AND METHODS

### Ethical approval

All animal procedures were reviewed and approved by Florida State University (FSU) Laboratory Animal Resources (Protocol 202000045) that abided by the American Veterinary Medical Association (AVMA) and the National Institutes of Health (NIH). In preparation for OB slice electrophysiology, mice were anesthetized with isoflurane (Aerrane; Baxter, Deerfield, IL, USA) using the Institutional Animal Care and Use Committee (IACUC)-approved drop method and then were killed by decapitation (AVMA Guidelines on Euthanasia, June 2007). In preparation for histology, mice were anaesthetized with a mixture of ketamine (100 mg/kg body weight) and xylazine (10 mg/kg body weight) prior to intracardial perfusion with 4% paraformaldehyde and euthanasia by decapitation (AVMA Guidelines on Euthanasia, June 2007). Our work was guided by the animal ethics checklist reported by Grundy, 2015 (Grundy, 2015).

### Solutions and reagents

Salts and sugars, unless otherwise noted, were obtained from Thermo Fisher Scientific (Waltham, MA, USA) or Sigma-Aldrich Chemical (St. Louis, MO, USA). Artificial cerebral spinal fluid (aCSF) contained (in mm): 119 NaCl, 26.2 NaHCO_3_, 1 NaH_2_PO_4_, 2.5 KCl, 1.3 MgCl_2_, 2.5 CaCl_2_ and 22 d-glucose (pH 7.3; 300–310 mOsm). Sucrose-modified aCSF was used for vibratome sectioning and recovery, and contained (in mm): 83 NaCl, 26.2 NaHCO_3_, 1 NaH_2_PO_4_, 3.3 MgCl_2_, 0.5 CaCl_2_, 72 sucrose and 22 d-glucose (pH 7.3; 300-310 mOsm) (Jan and Westbrook, 2007). Slice intracellular (pipette) solution contained (in mm): 135 potassium gluconate, 10 KCl, 10 HEPES (N-2-hydroxyethylpiperazine-N-2-ethane sulfonic acid), 10 MgCl_2_, 0.4 NaGTP and 2 NaATP (pH 7.3; 280–285 mOsm). PBS contained the following (in mm): 136.9 NaCl, 2.7 KCl, 10.1 Na_2_HPO_4_, and 1.8 KH_2_PO_4_ (pH 7.4). Paraformaldehyde (PFA) for tissue fixation consisted of 4% PFA (Eastman Kodak, Rochester, NY, USA) in PBS. Blocking buffer and incubation buffer consisted of 2% bovine serum albumin fraction V (BSA) (Invitrogen, Waltham, MA, USA) and 0.2% Tween-20 (Fisher, Hampton, NH, USA) in 1X PBS (Salameh et al., 2017).

The following synaptic blockers and channel vestibule blocker were prepared as concentrated stock solutions in aCSF, stored at -20°C until day of use, and then diluted to the following working concentrations: 10 μM gabazine (catalog #104104-50-9, MilliporeSigma, Burlington, MA, USA), 5 μM 2,3-dioxo-6-nitro-7-sulfamoyl-benzo[f]quinoxaline (NBQX, catalog #ab120045, Abcam, Waltham, MA, USA), 25 μM (2R)-amino-5-phosphonovaleric acid (AP5, catalog #ab120003, Abcam), 1 nM margatoxin (MgTx, catalog # M8278, MilliporeSigma), 0.1 uM insulin (Fadool et al., 2011*a*) (catalog #I2643, MilliporeSigma), and 100 nM tetrodotoxin (TTX, catalog # 120055, Abcam). Prior to use, MgTx and insulin were additionally mixed with 0.05% bovine serum albumin (BSA fraction V, catalog # BSAV-RO, Sigma-Aldrich) to prevent loss of peptide to tubing and plasticware while recording.

### Antibodies

Monoclonal antibodies targeting the beta subunit of the IR or green fluorescent protein were purchased from Santa Cruz Biotechnology (Dallas, TX, USA). Anti-IR antibody was purchased pre-conjugated to Alexa Fluor 647 (Anti-INSR catalog # sc-57342 AF647; RRID:AB_784102); validated by Wu and associates (Wu et al., 2019). Anti-IR antibody was generated against the C-terminus of the beta subunit of human insulin receptor protein. FSU120, a rabbit polyclonal antiserum, was generated against the 46-amino acid sequence 478-MVIEEGGMNHSAFPQTPFKTGNSTATCTTNNNPNDCVNIKKIFTDV-523, representing the unique coding region of the Kv1.3 channel between transmembrane domain 6 and the carboxy terminal (Fadool *et al*., 2000). The purified peptide was produced by Genmed Synthesis (San Francisco, CA, USA), the antiserum was produced and then affinity purified by Cocalico Biologicals (Reamstown, PA, USA). We have validated this epitope in several previous studies (Fadool et al., 2000, 2004; Colley et al., 2007; Marks and Fadool, 2007; Biju et al., 2008; Marks et al., 2009) and this latest regeneration of the antisera, specifically (Velez et al., 2016; Kolling et al., 2022). Alexa Fluor 546 (Invitrogen, catalog #A-11035; RRID:AB_2534077) was used to label the FSU120 antibody for imaging.

### Animals

All mice were housed at the FSU vivarium on a standard 12 h/12 h light/dark cycle and in accordance with institutional requirements for animal care. Mice were individually housed in conventional-style rodent cages containing separate food and water that could be obtained *ad libitum*. Mice were maintained on chow that was comprised of 13.5% kcal fat, 59.1% kcal carbohydrate, and 28.05% kcal protein (Catalog #5001, Purina Company, St. Louis, MO, USA).

B6:CBA-Tg(Tbx21-cre)1Dlc/J mice with cre recombinase expression limited to mitral and tufted cells (catalog #024507, Jackson Labs, Bar Harbor, ME, USA; RRID:IMSR_JAX:024507) were crossed with B6J.129(B6N)-Gt(ROSA)26Sor^tm1(CAG-cas9^,-EGFP)Fezh^/J mice (catalog #026175, Jackson Labs; RRID:IMSR_JAX:026175) to generate progeny mice with bicistronic spCas9/EGFP expression limited to the mitral and tufted cells.

The EGFP signal, along with anticipated physiological parameters, was used to identify mitral vs. tufted cells (Chen and Shepherd, 1997; Marks et al., 2009). A total of 78 Tbx21-cre x Cas9/EGFP male and female mice were used in our experiments for electrophysiology and RNAscope (referred to as simply ‘GFP mice’ throughout) and ranged from 19 to 28 postnatal days (PD) in age. For Tbx21-cre x Cas9/EGFP mice that were used for immunocytochemistry, mice were virally infected with an adenovirus carrying a sgRNA [GCT GCC GCC AGA CAT GAC CG] that targeted a protospacer adjacent motif (PAM recognition) target site that had close proximity to the ‘start’ codon of Kv1.3 to achieve CRISPR editing of the channel selectively from mitral and tufted cells. sgRNA was packaged into AAV9 viral particles (∼3E+12 VG/ml) using a U6 promotor design (AAV9-hSyn-mCherry-U6-sgRNA virus particle; SignaGen Laboratories). Retroorbital injection of the sgRNA virus was performed on neonatal Tbx21-Cre x Cas9/EGFP mice at age PD5 as previously described (Kolling et al., 2022). Immunocytochemistry experiments were performed in mice that were subsequently raised to adult (greater than 6 month).

Kv1.3-/- mice were crossed with Thy1-YFP mice to generate progeny mice with fluorescently-labeled OB neurons. Kv1.3-/- mice (RRID:IMSR_JAX:027392) were produced previously by excision of the N-terminal one-third of the *Kv1.3* gene (Xu et al., 2003; Koni et al., 2003) and were a generous gift from Drs. Leonard Kaczmarek and Richard Flavel (Yale University, New Haven, CT, USA). Thy1-YFP mice (RRID:IMSR_JAX:014130) were a gift from Dr. Gouping Feng (MIT Broad Institute, Boston, MA, USA) (Feng et al., 2000). A total of 5 Kv1.3-/- x YFP male and female progeny mice were used for electrophysiology (referred to as simply ‘Kv1.3-/- mice’ throughout) and ranged from PD19-28 in age. Homozygous Kv1.3-/- mice (not crossed with Thy1-YFP) were also used for immunocytochemistry experiments and compared with wildtype C57BL6/J lines (RRID:IMSR_JAX:000664) to examine specificity of the Kv1.3 channel plus IR kinase labeling.

### Olfactory bulb slice recordings

OB slices for electrophysiological recordings were acquired from GFP or Kv1.3-/- mice as described previously (Fadool et al., 2011; Mast and Fadool, 2012; Huang et al., 2017; Thiebaud et al., 2019; Kolling et al., 2022). Briefly, mice were anesthetized by inhalation of isoflurane, quickly decapitated, and the OBs were rapidly exposed by removing the dorsal and lateral portions of the skull (Nickell et al., 1996). The OBs (while still attached to the forebrain) were quickly removed, glued to a sectioning block with Superglue (Lowe’s Home Improvement, Tallahassee, FL, USA) and submerged in oxygenated (95% O_2_/5%CO_2_), ice-cold, sucrose-modified artificial cerebral spinal fluid (aCSF) to prepare the tissue for sectioning. Coronal sections (300 μm) were cut in oxygenated, ice-cold, sucrose-modified aCSF using a Series 1000 Vibratome (Leica, Wetzlar, Germany; RRID:SCR_016495). The OB sections were allowed to recover in an interface chamber (Krimer and Goldman-Rakic, 1997) with oxygenated, sucrose-modified aCSF at 33° C for 15 min and then were maintained at room temperature (rt) in oxygenated normal aCSF until needed (Fadool *et al*., 2011). OB slices were recorded in a continuously-perfused (Ismatec; 1-2 ml/min), submerged-slice recording chamber (RC-26, Warner Instruments, Hamden, CT, USA) with aCSF at rt.

OB slices were visualized at 10x and 40x using an Axioskop 2FS Plus Microscope (Carl Zeiss Microimaging, Thornwood, NY, USA) equipped with fluorescent excitation and infrared detection capabilities (CCD100; Dage-MTI, Michigan City, IN, USA). Electrodes were fabricated from borosilicate glass (#1405002; Hilgenberg GmbH, Malsfeld, Germany) to a diameter of ∼2 μm to yield pipette resistances ranging from 4 to 8 MΩ. Positive pressure was retained when navigating through the OB laminae until a high resistance seal (1.5–10 GΩ) was obtained on a positionally-identified MC or sTC in the slice (Fadool et al., 2011). The morphology and biophysical properties of the neurons were used to distinguish MCs and sTCs (Nickell et al., 1996; Schoppa and Westbrook, 2001; Fadool et al., 2004; Fadool and Kolling, 2020). MCs were selected only if > 50% of their cell body resided within the MCL. sTCs were selected only if their cell body resided in the outer one-third of the EPL, and were not selected if their cell body partially resided within the GLM. In addition, GFP mice provided fluorescent markers that served as secondary confirmation of cell identity, and were readily distinguishable. The cell-attached configuration was established by applying gentle suction to the lumen of the pipette while monitoring resistance. Whole-cell access was obtained by applying a 100 ms, 1 mV pulse across the membrane while applying sharp suction to the lumen of the pipette. Membrane voltage and current properties were generated using pCLAMP, version 9 or 10, in conjunction with a Multiclamp 700B amplifier (Axon Instruments; RRID:SCR_011323/Molecular Devices; RRID:SCR_018455). The analog signal was filtered at 3 kHz and minimally digitally sampled every 100 μs with a Digidata 1440A digitizer (Axon Instruments, Molecular Devices; RRID:SCR_021038). The pipette capacitance was electrically compensated through the capacitance neutralization circuit of the Multiclamp 700B amplifier. Resting membrane potentials were corrected for a calculated -7 mV junction potential offset. Membrane capacitance was acquired from the membrane test function of Clampex 10.7 (Axon Instruments, RRID:SCR_011323). Input resistance was calculated using Ohm’s law by injecting a -50 pA current and measuring the induced voltage deflection. For clarity across disciplines, this report uses the electrophysiological terminology that is used by the pCLAMP software suite. Where applicable, other common names are included in parentheses.

After establishing a whole-cell configuration, cells were first sampled for adequate resting potential (less than -50 mV), proper access resistance (less than 40 MΩ), and adequate input resistance (greater than 150 MΩ) prior to initiating either a series of current- or voltage-clamp recordings. For current-clamp recordings, cells were held at -60 mV (near rest; V_h_) between pulses to compare data across cells. Peri-threshold current levels were determined by incrementally injecting a family of current pulses of 1000 millisecond (ms) duration (P_d_). Current was injected every 30 seconds (s) and was stepped in 25 pA increments from -50 pA to 175 pA to obtain action potential (AP) characteristics and current-dependence data. Following the determination of spike threshold, cells were stimulated with 10 super-threshold sweeps using a P_d_ of 5000 ms (typically ranging from 15 to 150 pA) every 30 s (T_s_) to acquire spike frequency and latency data. Current-clamp data were acquired in the presence of synaptic blockers and voltage-clamp data were acquired in the presence of 20 μM TTX to eliminate unwanted AP spiking (see Solutions).

AP characteristics, latency to first spike, spike event frequency, interspike interval, intratrain interval, and spike train length were measured using Clampfit software as described previously (Balu et al., 2004; Fadool et al., 2011; Mast and Fadool, 2012). A train was defined as three or more consecutive spikes within a period of 100 ms or less, as established by Balu et al. (Balu et al., 2004). M/TC firing is intrinsically intermittent and is characterized by variable train characteristics. As such, classical means of computing spike timing variability, such as peri-stimulus time histograms, were less suitable for the behavior of these neurons. Therefore, alternative means of spike analysis were applied as described previously (Balu et al., 2004; Fadool et al., 2011; Thiebaud et al., 2016; Kolling et al., 2022). For voltage-clamp recordings, cells were held at -80 mV (V_h_), then stepped from -100 mV to +40 mV (V_c_) in +20 mV increments for a duration of 400 ms (P_d_) with a stimulation interval (T_s_) of 45 s.

### Tissue Preparation and Immunocytochemistry

Mice were deeply anesthetized and then terminated by cardiac perfusion using 4% PFA as previously described (Thiebaud et al., 2014). The mice were decapitated, and brains carefully removed from the skulls to flash freeze the brain in protective medium (Biju et al., 2008). More detailed immunocytochemistry procedures can be found in our previous works (Biju et al., 2008; Thiebaud et al., 2014), and current parameters for this project were as follows. Coronal sections at 25-µm thickness (Model 1850UV, Leica Biosystems) were prepared for slide-mounting. Labeling control consisted of blocking and incubation in buffer without antibodies. No-primary (omission of FSU120 primary antisera) and singly-labeled controls were performed to determine the specificity of the anti-rabbit secondary antibody with pre-conjugated antibodies. Sections were blocked with 3% goat sera and incubation of primary antisera took place overnight at 4°C (1:1000 anti-INSR, 1:1000 FSU120) with secondary antisera (1:500 Alexa Fluor 486) incubation following the next day for 2 h at room temperature. Slides were cover-slipped using Vectashield mounting media containing DAPI (H1200-10, Vector Laboratories, Newark, CA) before imaging. Imaging was performed on a Yokogawa Series Nikon Spinning Disk Confocal Microscope (Nikon Instruments, Inc., Melville NY, USA; Model CSUW1, RRID:SCR_021741) using a 20X lens, and appropriate laser lines for each reporter (475 nm, 551 nm, 642 nm). Post-hoc analyses were performed using Photoshop CS4 (Adobe, San Jose, CA, USA; RRID:SCR_014199). The minimum and maximum thresholds for each color channel were adjusted uniformly for all images before using the “difference” feature to align multidimensional images in Photoshop. This method allowed for transient removal of specific channels in order to more clearly determine co-localization of reporters.

### RNA scope and quantification

Five male and five female GFP mice (10 mice total) were anesthetized with ketamine/xylazine and fixed via transcardial perfusion of PBS followed by 4% PFA. Brains were cryoprotected in 30% sucrose before being embedded in OCT and flash frozen. Mouse brains were cryosectioned coronally at 16 µm, and sections were collected 1:10. Sections were mounted on histobond glass slides, and RNAscope was performed according to the ACDbio manufacturer’s protocol using commercially available probes for *GFP, Kv1.3* and *Insr*. The sequence of the mouse *GFP* transgene was used to verify the selection of GFP RNAscope probe. Images were acquired using an Olympus VS200 slidescanner at 20X, and analysis was performed using QuPath software (v 0.5). The ‘cell detection’ function was first used to identify Tbx21+ cells, then object classifiers were created for *Kv1.3* and *Insr* using a threshold for mean fluorescent intensity within each cell. Validation of this workflow was performed manually, once per neurolaminar layer (MCL, EPL, and GLM).

### Experimental design and statistical analyses

Before performing any statistical comparisons, data were first analyzed with the Dixon’s Q test to identify any outliers. Data were then checked for normal distribution and homogeneity of variance using the Fmax test. Analysis of data collected did not identify any outliers. Where collected data violated homogeneity of variance (failed the Fmax test), the selection of applied statistics was changed to the nonparametric equivalent. Paired comparisons (innate properties, action potential kinetics, bursting metrics) were analyzed using Student’s *t* test or Mann-Whitney U at the 95% confidence level (α < 0.05). There were no sex-specific differences in the measured parameters, so measurements from both sexes were combined. IV plots, frequency and interspike interval (ISI) were analyzed using a mixed two-way RM ANOVA with current and voltage as factors (IV) or current and cell type as factors (frequency and ISI); at the 95% confidence level (α < 0.05). Modulation of voltage-activated current between cell types was analyzed using a mixed three-way RM ANOVA with voltage, cell type, and drug as factors; at the 95% confidence level (α < 0.05). Insulin modulation data were analyzed using within-cell comparisons by applying a paired *t* test at the 95% confidence level (α < 0.05). Categorical AP spike train type analysis was performed using a Chi-Square test, where the expected proportion of sTCs eliciting spike trains was set as the observed proportion of MCs eliciting spike trains; 95% confidence level (α < 0.05). For the two-way RM ANOVA tests, the Sidak method for multiple comparison testing was used as the post-hoc analysis to make mean-wise comparisons between cell types or between drug treatments. All reported values in the text represent the mean ± SD. Figures largely report individual data points using violin graphs and represent number of cells as noted in the figure legends.

Sample sizes for the line graphs are reported in the figure legends. Where means are reported in the figures, the error is reported as the SEM. Individual *p* values are reported for each experiment within the corresponding graph with the individual F statistic described in the results section. Statistical tests were performed using GraphPad Prism 9 (GraphPad Software) while the graphs and figures were produced using a combination of Origin v8 (OriginLab Corporation; RRID:SCR_014212), Adobe Photoshop CS4 (Adobe; RRID:SCR_014199), and GraphPad Prism 9 (GraphPad Software; RRID:SCR_002798).

All electrophysiological data were analyzed using pClamp, version 10 (Clampfit 10.7; Molecular Devices; RRID:SCR_011323) and Igor Pro, version 6.12A (WaveMetrics Inc.; RRID:SCR_000325) with the plug-in NeuroMatic, version 2 (written by Jason Rothman; http://www.neuromatic.thinkrandom.com; RRID:SCR_004186). Exact *p* values, statistical tests, and parameters are given in the results text/figures for each data set. Specific variables required for independent replication (sample size, number of animals, etc.) and full statistical reporting can be found in the methods and results text/figures for each respective experiment and the supplemental statistical table (Supplemental Table 1).

## ACKNOWLEDGEMENTS

We would like to thank Carley Huffstetler, Alexis Cox, Tyla Dolezel, and Franklin Pacheco for routine technical assistance and in maintaining mouse colonies. We would like to thank Jaime White-James for the oversight of our animal husbandry and routine mouse care needs.

## AI USE

The authors declare that no AI tools were used for this manuscript including scientific writing, generation of code, production of images, or in the collection or analyses of data.

## AUTHOR CONTRIBUTIONS

All authors critically evaluated the work for important intellectual content. All authors acquired, analyzed, or interpreted data in the study. As such, we confirm that all approved the final version of the manuscript; agreed to be accountable for all aspects of the work in ensuring that questions related to the accuracy or integrity of any part of the work are appropriately investigated and resolved; and all persons designated as authors qualify for authorship, and all who qualify for authorship are listed. In using the CRediT (Contributor Roles Taxonomy) guidelines, L.K. and D.F. were responsible for project conceptualization, data curation, formal analysis, funding acquisition, methodology, validation, visualization, and writing the original draft, reviewing, and editing. L.K. was responsible for electrophysiological recordings. S.M. was responsible for methodology and immunocytochemistry experiments. C.M. and D.F. were responsible for project administration and supervision. L.K., C.M. and D.F. were responsible for the provision of resources.

## DATA AVAILABILITY

All data referenced in this report are summarized in the primary figures contained in the publication. All summarized electrophysiological recordings data are stored in the Fadool Laboratory and are available upon request.

